# Rate dependent Effects of AMPH on attentional processes as measured by 5C-CPT task

**DOI:** 10.64898/2026.10.04.756500

**Authors:** Sara Abdulkader, John Gigg, Joanna C. Neill

**Affiliations:** University of Manchester

## Abstract

**Background:** ADHD is characterized by inattention and impulsivity. Stimulants such as amphetamine are the first line treatments for ADHD. Amphetamine (AMPH) works by increasing extracellular levels of dopamine and noradrenaline. Such stimulants are used to test predictive validity of animal tasks, where it has been reported that amphetamine improves rat 5-CSRTT accuracy when the session length is increased to 250 trials. This improvement in accuracy was observed later in the session as control performance declined. Proving that amphetamine can improve 5C-CPT task performance adds to the predictive validity of the task. No previous works has demonstrated the pro-attentional effects of amphetamine on sustained attention in rat 5C-CPT. We have previously shown that pharmacological manipulations of dopaminergic system improve waiting impulsivity in a rate dependent manner and the importance of analysing the data using Oldham method. The present study extends this work to investigate the effect of amphetamine on attentional processes.

**Objective:** To investigate effects of amphetamine on attentional processes as measured by 5C-CPT task using correlational analysis and group performance.

**Methods:** Trained female Lister hooded rats were tested with AMPH (0.1, 0.2 and 0.4 mg/kg) under reduced stimulus duration schedule (0.25,0.5,1,1.5 and 2s SD) over normal length session (120 trials). Correlational analysis (Oldham method) was used to understand the relationship between baseline performance measures and the change following administration of AMPH.

**Results:** The highest dose of AMPH impaired attentional performance in the 5C-CPT under reduced SD challenge (0.5s,1s,1.5s and 2s). This impairment was due to increase in FAR. Previous studies in 5-CSRTT suggest that reducing the SD increases attentional demand in the form of reducing accuracy without affecting waiting impulsivity. However, the 5-CSRTT does not include non-target trials, therefore, it cannot be used to measure sustained attention (SI) and responsivity index. The 5C-CPT requires greater attentional resources than the 5-CSRTT.This study is the first to report that reducing the SD changes responsivity index (bias) rather than measures of sustained attention (sensitivity index) in the 5C-CPT. Control sensitivity index did not change until the SD was reduced to less than 0.5s and this change was due to a change in baseline responsivity index (bias). Performance under each reduced SD revealed that the highest dose of AMPH (0.4 mg/kg) showed a tendency toward reducing sensitivity index at 0.5, 1.0 and 1.5s SDs. Under 1.0s SD, the highest dose of AMPH changed sustained attention (SI, HR and FAR) and omission in a rate-dependent manner without changing responsivity index. Performance across the session revealed that the impairment in sustained attention was observed at the beginning of the session. In the third trial block, Oldham method revealed an inverse relationship between baseline accuracy and the change following administration of 0.2 mg/kg AMPH, indicative of a rate-dependent effect.

**Conclusions:** This study successfully demonstrates rate-dependent effects of AMPH on measures of attentional performance in the 5C-CPT. This experiment also demonstrates that responsivity index is sensitive to reduced SD challenge. To our knowledge this is the first study to report the presence of a direct relationship between baseline sustained attention and magnitude of improvement following administration of 0.4 mg/kg AMPH and an inverse relationship between baseline accuracy and the change following administration of 0.2 mg/kg AMPH in 5C-CPT task. Future studies should test lower doses of amphetamine (0.05 and 0.1 mg/kg) over extended session length (250 trials) to reveal the pro-attentional effects of this drug on 5C-CPT performance measures.

## Introduction

Attention deficit hyperactivity disorder (ADHD) is a psychiatric disorder characterized by three main behavioural deficits: lack of attention, hyperactivity and impulsivity (Mueller et al., 2017). These are observed during early childhood and often persist into adult life. The prevalence of ADHD is between 2-7%: 5% of children have difficulties with inattention and hyperactivity and 2-4% of adults are diagnosed with ADHD (Sayal et al., 2018). With increasing age, inattention becomes more prominent while impulsivity diminishes (Faraone et al., 2015; Telzer et al., 2016). Children diagnosed with ADHD have a higher risk of developing social and cognitive problems in later life (Lara et al., 2009). Depending on the main symptoms, ADHD can be divided into three categories: lack of attention (most prevalent among girls), hyperactivity (more prevalent among boys) and a third subtype that is a combination of the first two (Rasmussen and Gillberg, 2000). ADHD is more common in boys than in girls. However, ADHD is unrecognised and underdiagnosed in many countries. In the UK less than 50% of children with inattention and hyperactivity have been diagnosed with ADHD (Faraone et al., 2015). Stimulants such as methylphenidate and amphetamine are the most common pharmacological treatment for ADHD (Mueller et al., 2017).). However, stimulants have the potential to lead to abuse and dependence (Ahn et al., 2013). For example, stimulants (smart pills) are commonly used by students to enhance academic performance (Sharma et al., 2014). Non-stimulants are the second line treatment for ADHD; however, they are less effective than stimulants and they have a delayed onset of action. Therefore, there is need for new therapeutic agents without the dependence potential (Sharma et al., 2014).

Dysfunction in dopaminergic system have been implicated in ADHD (Faraone et al., 2015). Stimulants are used in preclinical studies to test the predictive validity of tasks; for example, methylphenidate, which improves inattention and impulsivity in human subjects, has been shown to improve 5C-CPT performance of low-attentive and high-impulsive rats under challenging task conditions (Tomlinson et al., 2014) demonstrating the predictive validity of 5C-CPT task. Amphetamine is commonly used to treat ADHD, however, despite its positive clinical effects, few preclinical studies have reported pro-attentional effects in rats. In 5-CSRTT, low doses of amphetamine improved attentional accuracy (Grottick and Higgins, 2002). Clinical studies suggest that stimulants improve performance in a baseline-dependent manner (Allman et al., 2010). It has been reported that correlational analysis using Oldham’s method can help in revealing baseline-dependent effects of stimulants on attention and impulsivity (Bickel et al., 2016). Tomlinson et al. (2014) divided rats into two sub-groups based on their baseline performance. The highest dose of methylphenidate improved accuracy of low-attentive rats without having a significant effect on high-attentive rats. This finding suggests the presence of an inverse relationship between baseline accuracy and the drug-induced change; however, correlational analysis was not used in this study. Sensitivity index represents the ability of rats to distinguish between target and non-target trials. In the same study, the other two doses of methylphenidate (0.5 and 1 mg/kg) improved the sensitivity index (sustained attention) of low-attentive rats and showed a tendency toward reducing sensitivity index of high-attentive rats (Tomlinson et al., 2014). However, this improvement was due to a change in responsivity index rather than attentional processes (Unpublished data). In other words, 0.5mg/kg methylphenidate significantly decreased hit rate and showed a tendency toward reducing false alarm rate. Together, these effects contributed to a decrease in responsivity index from an average of (-0.09 ± 0.07) to (-0.15 ± 0.11). This finding suggest that the predictive validity of 5C-CPT tasks is yet to be determined.

In the signal detection task, low doses of amphetamine (0.1mg/kg and 0.25 mg/kg) improved attentional accuracy in a rate-dependent manner when rats were tested under conditions that required greater attentional resources (reduced SD challenge) (Turner et al., 2016). The signal detection task is different from 5C-CPT in that the animal is required to detect a target visual stimulus presented repeatedly in the same location (central panel) rather than detecting a stimulus that may appear in one of five locations (Turner et al., 2016). Correlational analysis revealed a moderate positive association between baseline performance and the change following administration of 0.1 or 0.25 mg/kg amphetamine in signal detection task (Turner et al., 2016). The rate-dependent effect was consistent with a full range effect of baseline accuracy in which low baseline accuracy values increased while high baseline accuracy values decreased following administration of low doses of amphetamine. We previously reported that amphetamine improves waiting impulsivity in a baseline dependent manner when rats were tested under extended ITI. It would be interesting to test whether amphetamine can improve attention in the 5C-CPT in a rate-dependent manner under challenging task conditions that target attentional processes. Therefore, a reduced SD challenge was used here as this type of challenge increases attentional demand (Turner et al., 2017; Caballero-Puntiverio et al., 2018). The aim of this study was, therefore, to examine the effects of amphetamine (0.1,0.2 and 0.4mg/kg) under reduced SD challenge in the rat 5C-CPT and to determine whether amphetamine can change performance in a rate-dependent manner. Rather than dividing rats into subgroups of high and low performers, correlational analysis using simple and Oldham methods were used to understand the relationship between baseline performance measures and the drug-induced change.

## Materials and Methods

### Animals

Female Lister hooded rats (n=15; Charles River, UK) were housed in groups of 5 on a reversed 12h light cycle in a temperature (21 ± 2 °C) and humidity (55±5%) controlled environment (University of Manchester BSF facility). Rats weighed 230±10g upon arrival. They were allowed free access to food (Special Diet Services, UK) and water for one week prior to the beginning of training. Two days before training commenced food restriction was initiated to encourage their task engagement and this continued throughout training. Rats were maintained at approximately 90% of their free-feeding body weight (fed 10 g rat chow/rat/day) and were provided with free access to water. All procedures were conducted in accordance with the UK animals (Scientific Procedures) 1986 Act and local University ethical guidelines.

### Rat 5C-CPT Apparatus

Training took place in 8 operant chambers (Campden instruments Ltd, UK), each placed inside a ventilated and sound-attenuating box. The chambers were of aluminium construction 25cm × 25 cm. Each chamber had a curved wall containing nine apertures. For 5C-CPT training, four of the apertures were blocked by metal caps while the other five were left open (numbers 1, 3, 5, 7 and 9 were open). Each aperture had a light and an infrared beam crossing its entrance to record beam breaks following nose poke. A reward dispenser was located outside each chamber that automatically dispensed reinforcers into a food tray at the front of the chamber. Entrance to each food tray was covered by a hinged panel so that tray entries were recorded when the rat’s snout opened the panel. House lights were placed in the side wall of the chambers and performance was monitored by a camera attached to the roof.

### 5C-CPT behavioural training

Unlike 5-CSRTT, the 5C-CPT incorporates non-target trials in which all five holes are lit and the rat must not respond at any of the five reward positions (Young et al., 2009). The incorporation of non-target trials helps to dissociate response inhibition from waiting impulsivity; premature responses during target trials reflect a deficit in waiting impulsivity, while the failure to withhold responding (false alarms) during non-target trials represents response disinhibition (Young et al., 2011). This task also allows the assessment of vigilance in rodents in a manner consistent with human CPT using signal detection theory (Tomlinson et al., 2014). In addition, a variable ITI is incorporated in 5C-CPT to prevent rats from simply timing their response (Van Enkhuizen et al., 2013). 5C-CPT training was carried out 5 days per week. A session consisted of 120 trials or was 30 min long, whichever came first. The training consisted of 3 phases: (i) 5-CSRTT training, (ii) 5C-CPT training with fixed ITI and (iii) 5C-CPT training with variable ITI (Tomlinson et al., 2015). The task parameters such as stimulus duration and limited hold were gradually reduced for rats based on their individual performance. Some of the key parameters that are used to assess performance in the task are listed in Table 1.

**Table 1:** Description of 5-CPT outcome measures.

| **Outcome measures** | **Descriptions** | **Dimension** |
| --- | --- | --- |
| **Correct response (Hit)** | Response made in the illuminated hole |  |
| **Incorrect response** | Response made but in the wrong hole |  |
| **Error of Omission (Miss)** | Failure to respond to the stimulus within a certain period |  |
| **Accuracy** | Proportion of correct responses to incorrect responses | Selective attention |
| **Correct rejection** | Correctly withholding the response to non-target stimuli |  |
| **False alarm** | Failure to withhold the response to non-target stimuli |  |
| **Hit rate (HR)** | Proportion of 'hits' to 'misses' |  |
| **False alarm rate (FAR)** | Proportion of false alarms to correct rejections | Response inhibition |
| **Sensitivity index** | Ability of rats to distinguish between target and non-target trials | Sustained Attention |
| **Responsivity index (Bias)** | Tendency to respond to stimuli | Motivation |
| **Premature response** | Response made before stimulus presentation | Waiting impulsivity |
| **Perseverative response** | Repetitive responses (repeated nose pokes) even after making a response | Compulsive behaviour |
| **Correct latency** | Time taken to make a correct response | Attention |
| **Correct magazine** | Time taken to collect the reinforcer after the correct response is made | Motivation |

### Signal detection theory (SDT)

Two primary parameters are calculated from 5C-CPT trial-outcome measures. The first is hit rate, which is the proportion of ‘hits’ to ‘misses’ and the second is false alarm rate, which is the proportion of false alarms to correct rejections (Young et al., 2011; Table 2). From these two parameters, sensitivity (vigilance) and bias can be calculated using SDT (Young et al., 2011). Sensitivity reflects the ability to respond to the illuminated hole and inhibit that prepotent response when all five holes are illuminated (Young et al., 2009; Anderson, 2015). Sensitivity can be measured non-parametrically by calculating the sensitivity Index (SI). 5C-CPT performance can also be affected by bias (tendency to respond to stimuli), which can be measured non-parametrically by calculating the responsivity index (RI) (Young et al., 2013). A positive value for RI reflects bias towards FA (liberal response strategy), while a negative RI reflects bias towards omission (conservative response strategy). When RI is zero there is no bias (Anderson, 2015; Young et al., 2013).

**Table 2.** Parameters generated form 5C-CPT trials. Hit = correct response, Miss = omission, CR = correct rejection, FA = false alarm, FAR = false alarm rate, HR = hit rate, SI = sensitivity index and RI = responsivity index.

| Trial Type | Measure | Calculations | SDT parameters |
| --- | --- | --- | --- |
| Target trials | Hit and miss | $HR = \frac{Hit}{Hit + Miss}$ | $SI = \frac{HR - FAR}{2(HR + FAR) - (HR + FAR)^2}$ $RI = \frac{HR + FAR - 1}{1 - (FAR - HR)^2}$ |
| Non-target trials | CR and FA | $FAR = \frac{FA}{FA + CR}$ | |

### Experimental Design

#### Drug preparation

d-Amphetamine sulphate (Sigma, UK) was dissolved in 0.9% saline and injected via the intraperitoneal (i.p.) route at a volume of 1 ml/kg. All drug doses were calculated as base equivalent weight. The doses used in this study were chosen based on previous studies performed in rats (Barnes et al., 2012). AMPH doses were selected to be lower than those that increase locomotor activity (1.25 mg/kg) (Turner et al., 2016; 2017).

#### Behavioural testing

Fifteen rats were used to investigate the effect of amphetamine on 5C-CPT performance using challenging task conditions that consisted of reduced stimulus durations (0.25, 0.5, 1.0, 1.5 and 2.0s). Using a within-subjects design trained rats received 0.1, 0.2 or 0.4 mg/kg amphetamine or vehicle, 10 min before the task.

#### Statistical analysis

All data are displayed as observed mean ± SEM. Overall performance was analysed by one-way repeated measures ANOVA followed by Planned Comparisons. Analysis of the performance across the duration of the session was carried out by two-way repeated measures ANOVA (trial and treatment as within-subject factors) followed by Planned Comparison on the predicted means. If the outcome of the repeated-measures ANOVA yielded significant effects of dose, further post-hoc analysis was performed using Dunnett multiple comparison test. Analysis of the performance across the prolonged ITI was carried out by two-way repeated measures ANOVA (ITI and treatment as within-subject factors), followed by Planned Comparisons. Alpha level was set to 0.05 and all analysis was carried out using Prism (v8.0). One rat was excluded from the analysis due to large number of omissions.

#### Correlational analysis

Three methods are used in literature to assess the relationship between baseline and magnitude of improvement following drug administration (Table 3). This study investigated if baseline performance predicted the degree of change in performance following administration of GBR using Oldham method. Oldham’s method was used to determine the presence of a rate dependent relationship. Pearson’s correlation was used when individuals were normally distributed. Nonparametric Spearman’s correlation was used when individuals were not normally distributed. Oldham’s *r* > 0.3 indicates the presence of a rate dependent relationship (Oldham, 1962; Snider et al., 2016; Bickel et al., 2016).

**Table 3.** Methods used in literature to assess presence of a baseline dependent effect (Oldham, 1962; Bickel et al., 2016; Turner et al., 2017; Snider et al.,2018).

| Method | Descriptions |  |
| --- | --- | --- |
| Simple method | Correlation between baseline values and the drug induced change (Spurious correlation) | This method is affected by mathematical coupling and regression to the mean |
| Oldham's method | Correlation between average of pre- and post-treatment values and the drug induced change | Eliminates the effect of mathematical coupling and regression to the mean |
| Re-test method | Correlation between adjusted baseline value and the drug induced change | Reduces the bias associated with regression to the mean |

Regression to the mean is detected when repeated measurements are taken from the same subject. For example, if baseline premature response after vehicle administration is high, the probability of having a lower premature response after drug administration is greater than having a higher premature response and vice versa (Chiolero et al., 2013). Regression to the mean can also happen at sub-group levels (Chiolero et al., 2013). Mathematical coupling occurs during correlations where one variable is part of the calculation for the other variable, resulting in a spurious correlation (Tu et al., 2004). This false correlation will be present irrespective of any true relationship between the two measurements. In simple correlation, the baseline value is part of calculation of the change following drug administration (drug value – baseline value); therefore, these values are not independent as there is a mathematical coupling between them (Tu et al., 2004; Chiolero et al., 2013). This coupling induces a strong negative false correlation when baseline value is plotted against the change following drug administration. To avoid the biases associated with the simple method, one may assess the relationship between the average of pre- and post-treatment value with the change following drug administration (Oldham’s method). Here, Oldham suggested that the average is closer to the true baseline value than the single value obtained following vehicle administration (Oldham, 1962). In addition, the average is not part of calculation of the change (Tu et al., 2004; Chiolero et al., 2013). This method has been used in literature to determine the presence of a rate-dependent effect (e.g., Bickel et al., 2016; Snider et al., 2016; Snider et al., 2018).

## Results

### Effects of Amphetamine on 5C-CPT performance under reduced stimulus durations (0.25,0.5,1,1.5 and 2s)

The main effect of amphetamine on typical 5C-CPT performance indices is shown in Fig.1. One-way repeated measures ANOVA revealed no significant treatment effect on sensitivity index [F (2.607, 33.90) = 2.219, P=0.1114], responsivity index [F (2.837, 36.88) = 0.5699, P=0.6290], hit rate [F (2.621, 34.08) = 2.513, P=0.0821], false alarm rate [F (2.629, 34.18) = 0.06963, P=0.9649], accuracy [F (2.703, 35.13) = 1.175, P=0.3304] and omission [F (2.707, 35.19) = 2.560, P=0.0760], The highest dose of amphetamine showed a tendency towards reducing SI, reducing hit rate and increasing omissions.

**Figure 1:**
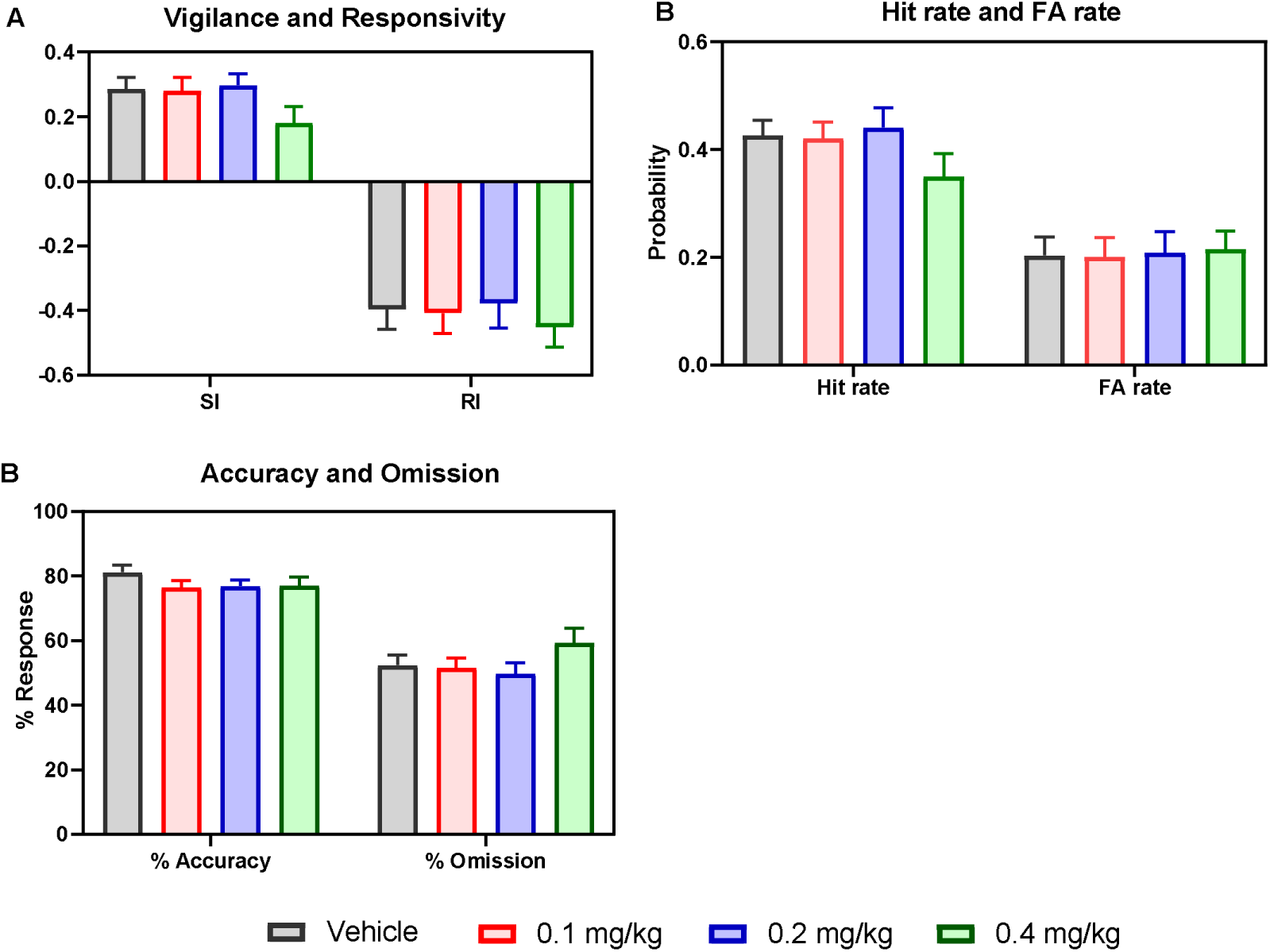
Effects of AMPH on 5C-CPT perfomance tested under reduced SD (n=14). (A) Vigilance and responsivity. (B) Hit rate and false alarm rate. (C) Accuracy and omission. Measures are shown as observed mean ±SEM.

The relationship between baseline 5C-CPT performance measures and the drug-induced change is shown in Fig. 6.2. Oldham method revealed insignificant associations between baseline SI, RI, HR, FAR or omission and AMPH induced change, indicating that AMPH does not modulate these measures in a baseline dependent manner. The relationships between baseline accuracy and the drug-induced change were affected by an outlier. Oldham method revealed a moderate positive and significant correlation between baseline accuracy and 0.4 mg/kg AMPH induced change after the outlier was excluded from the analysis, indicative of a rate-dependent effect (Fig 2).

**Figure 2:**
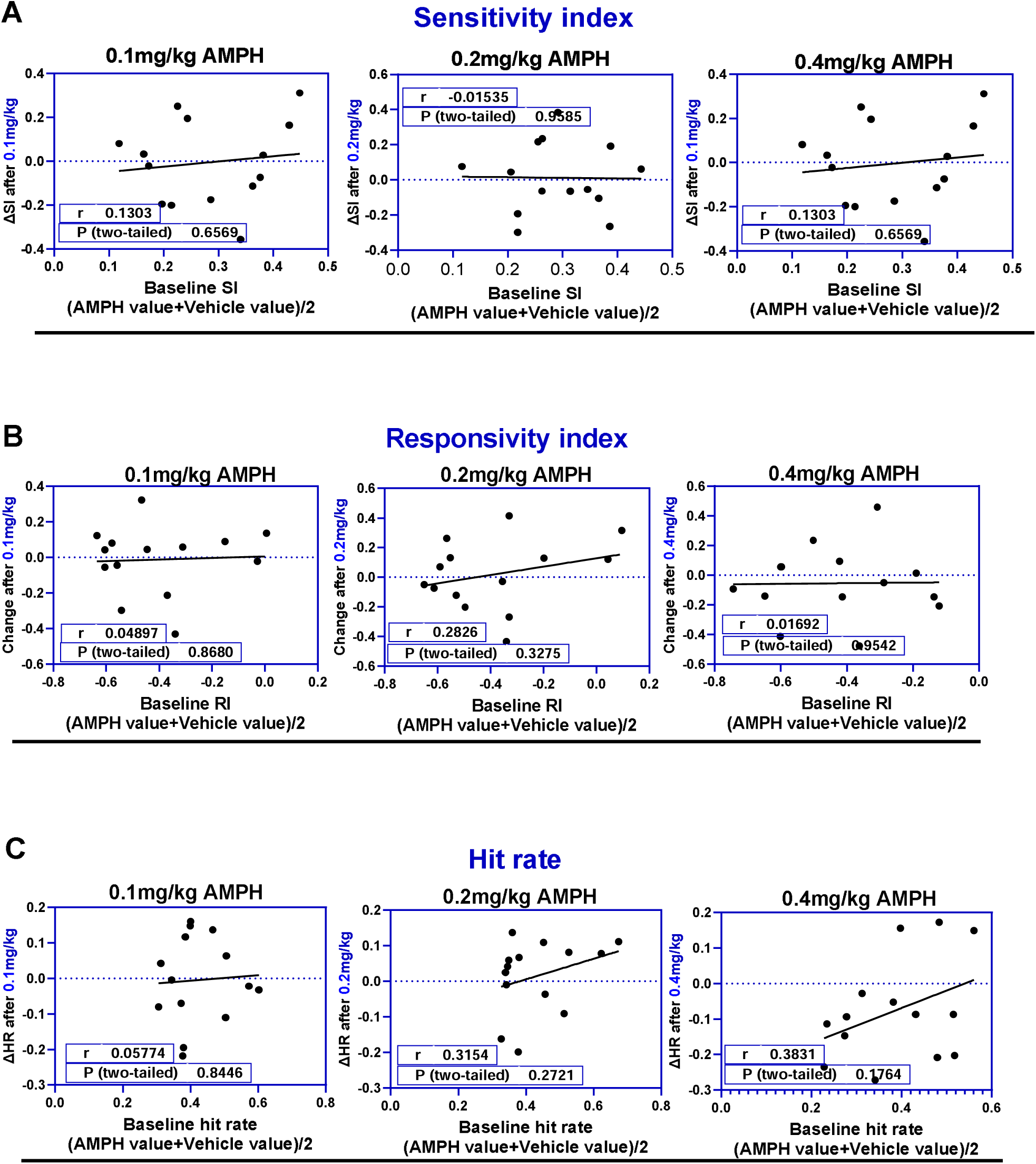
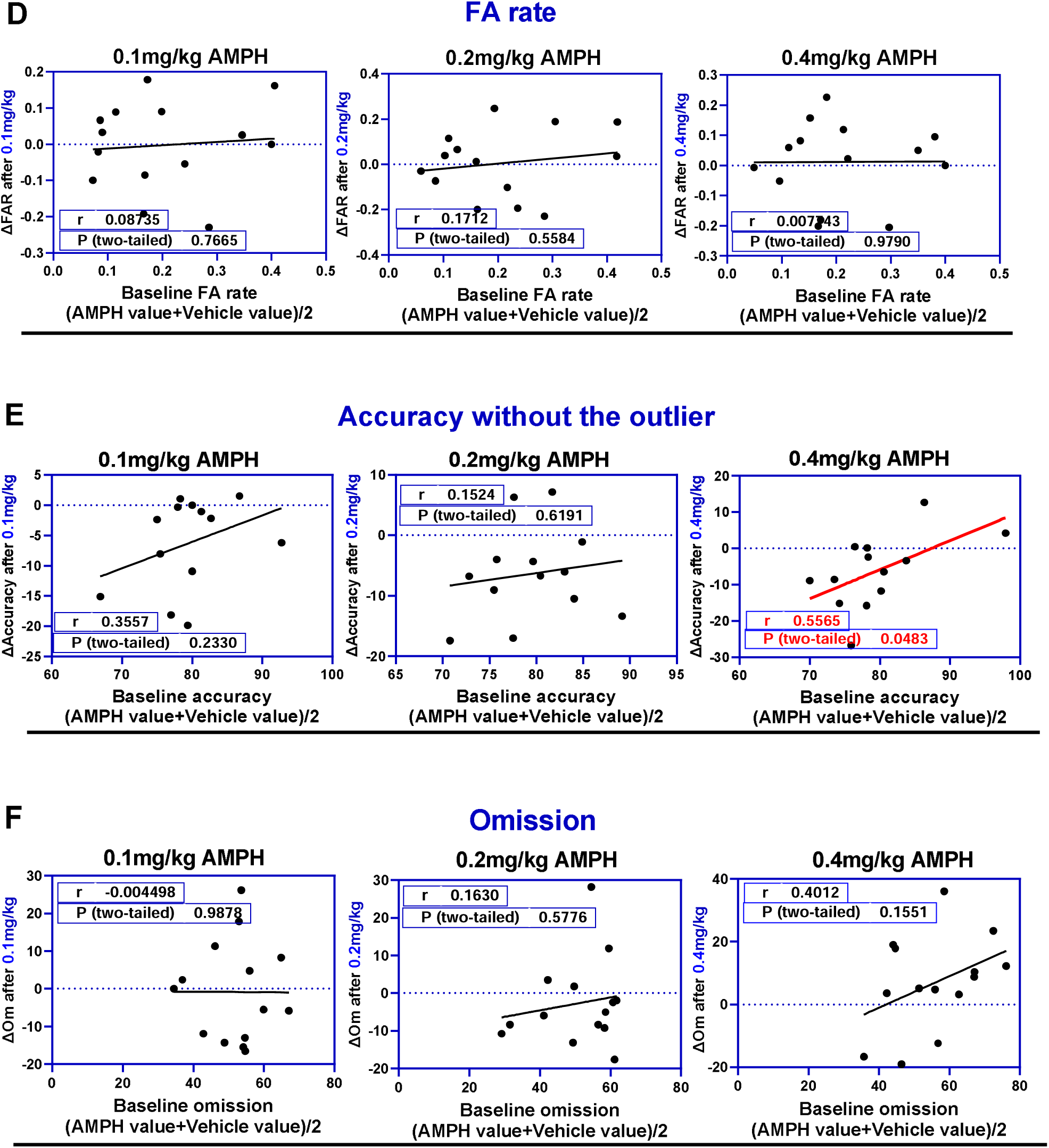
Relationship between baseline 5C-CPT performance measures and AMPH induced change using Oldham method (n=14). (A) Sensitivity index. (B) Responsivity index. (C) Hit rate. (D) False alarm rate. (E) Accuracy. (F) Omission. The red regression lines represent statistically the significant relationships.

#### Effects of Amphetamine on 5C-CPT performance under short SDs (0.5s,1s,1.5s and 2s SDs)

The number of omissions during the 0.25s SD was very high, representing a potential confound; therefore, we sought to test the effect on performance when the 0.25s SD was excluded from the analysis. A one-way repeated measures ANOVA revealed a significant treatment effect on SI. Dunnett’s multiple comparisons test revealed a significant decrease in SI following administration of the highest dose of AMPH compared with control (Fig. 3A). One-way repeated measures ANOVA revealed no significant treatment effect on responsivity index [F (2.862, 37.20) = 0.5333, P=0.6541], hit rate [F (2.441, 31.74) = 2.451, P=0.0924], false alarm rate [F (2.689, 34.96) = 0.6462, P=0.5743], accuracy [F (2.320, 30.1) = 2.847, P=0.0665] and omission [F (2.454, 31.90) = 2.529, P=0.0854].

**Figure 3:**
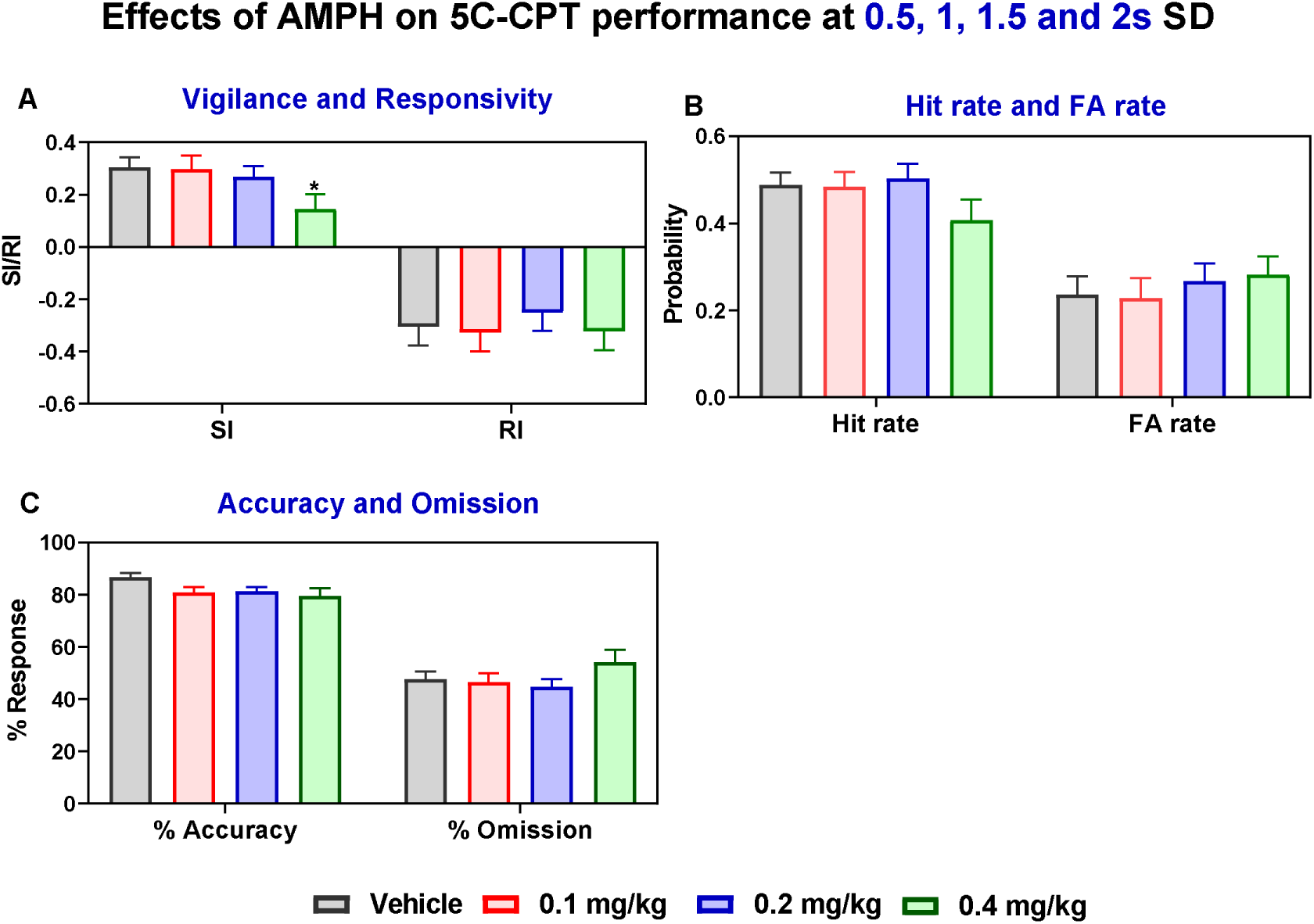
Effects of amphetamine on 5C-CPT performance of rats tested under reduced SDs (n=14). (A) SI and RI (B) Hit rate and false alarm rate. (C) Accuracy and omission. * *p* < 0.05 compared to Control (Dunnett). Measures are shown as observed mean ±SEM.

#### Effects of Amphetamine on 5C-CPT performance under short SDs (0.5,1,1.5)

In order to focus on the SDs that were most challenging for the rats we omitted two SDs; these were (a) 0.25s (as the number of omissions during 0.25s SD was very high) and (b) 2.0s (rats were very good at performing the task at this SD). Thus, a one-way repeated measures ANOVA on the SD range that only included 0.5, 1.0 and 1.5s revealed a significant treatment effect [F (2.638, 31.65) = 5.989, *p* < 0.01] on vigilance. Dunnett *post hoc* tests revealed a significant decrease in SI following administration of the highest dose of amphetamine *(p* < 0.01) compared to saline treated rats (Fig 4A). This impairment was due to a decrease in hit rate accompanied by an increase in false alarm rate. The highest dose of amphetamine also showed a tendency toward increasing omissions and reducing accuracy (Fig. 4C). One-way repeated measures ANOVA revealed no significant treatment effect on responsivity index [F (2.300, 29.89) = 0.1340, P=0.9000], hit rate [F (2.180, 28.34) = 3.231, P=0.0507], false alarm rate [F (2.587, 33.63) = 1.142, P=0.3414] and accuracy [F (2.728, 35.47) = 1.841, P=0.1618]. One-way repeated measures ANOVA revealed a significant treatment effect on omission [F (2.11, 27.46) = 3.561, *p* < 0.05] and a significant matching [F (13, 39) = 3.318, p < 0.01]. Dunnett *post hoc* tests revealed no significant change in omission following administration of the highest dose of amphetamine compared to vehicle treated rats.

**Figure 4:**
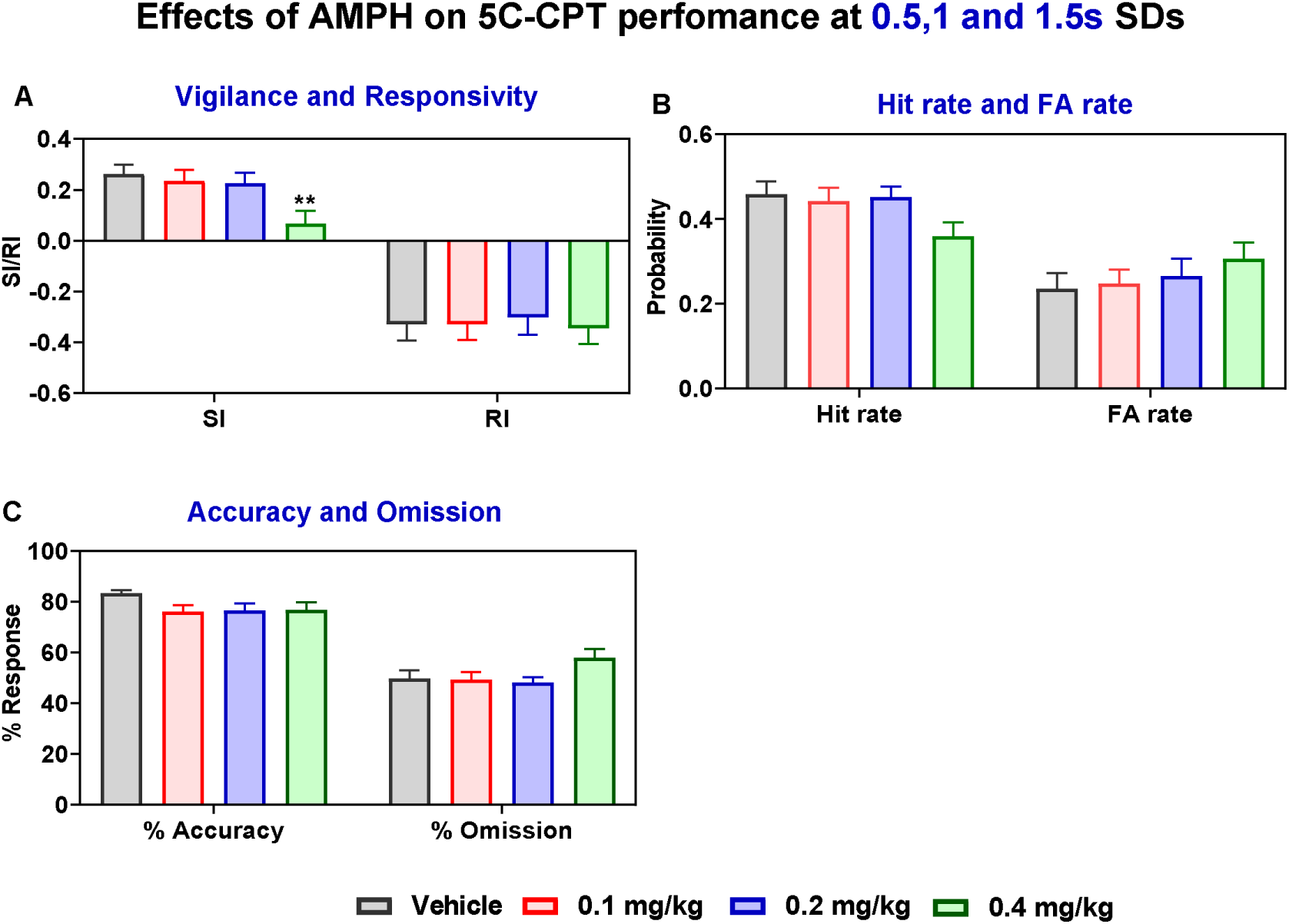
Differential effects of amphetamine on 5C-CPT performance measures of rats tested under short SDs (0.5,1,1.5s) (n=14). (A) *d* prime and responsivity index. (B) Hit rate and false alarm rate. (C) Accuracy and omission. ** *p* < 0.01 compared to Control (Dunnett). Measures are shown as observed mean ±SEM.

### Effects of amphetamine on 5C-CPT performance under each SD

#### Sensitivity index (Fig. 5A)

A two-way repeated measures ANOVA revealed a significant SD effect (F (3.003, 39.03) =8.634, *p* < 0.001) on sensitivity index and a significant treatment effect (F (2.610, 33.93) = 3.142, *p* < 0.05). Dunnett’s multiple comparison *post hoc* test revealed a significant increase in baseline sensitivity index at 0.5s (*p*< 0.01), 1.0s (*p*< 0.05) and 1.5s (*p*< 0.01) SDs when compared to saline treated rats at 0.25s, indicating that rats performed better at longer SDs. However, vehicle sensitivity index remained stable at 0.5, 1 and 1.5s SD despite the gradual increase in stimulus duration; impairment was only observed at 0.25s SD, suggesting that SI does not depend on SD length. The highest dose of amphetamine showed a tendency towards reducing sensitivity index at 0.25, 0.5s, 1.0s and 1.5s SD when compared to vehicle treated rats at the same SD. This impairment was less obvious at 2s SDs.

**Figure 5:**
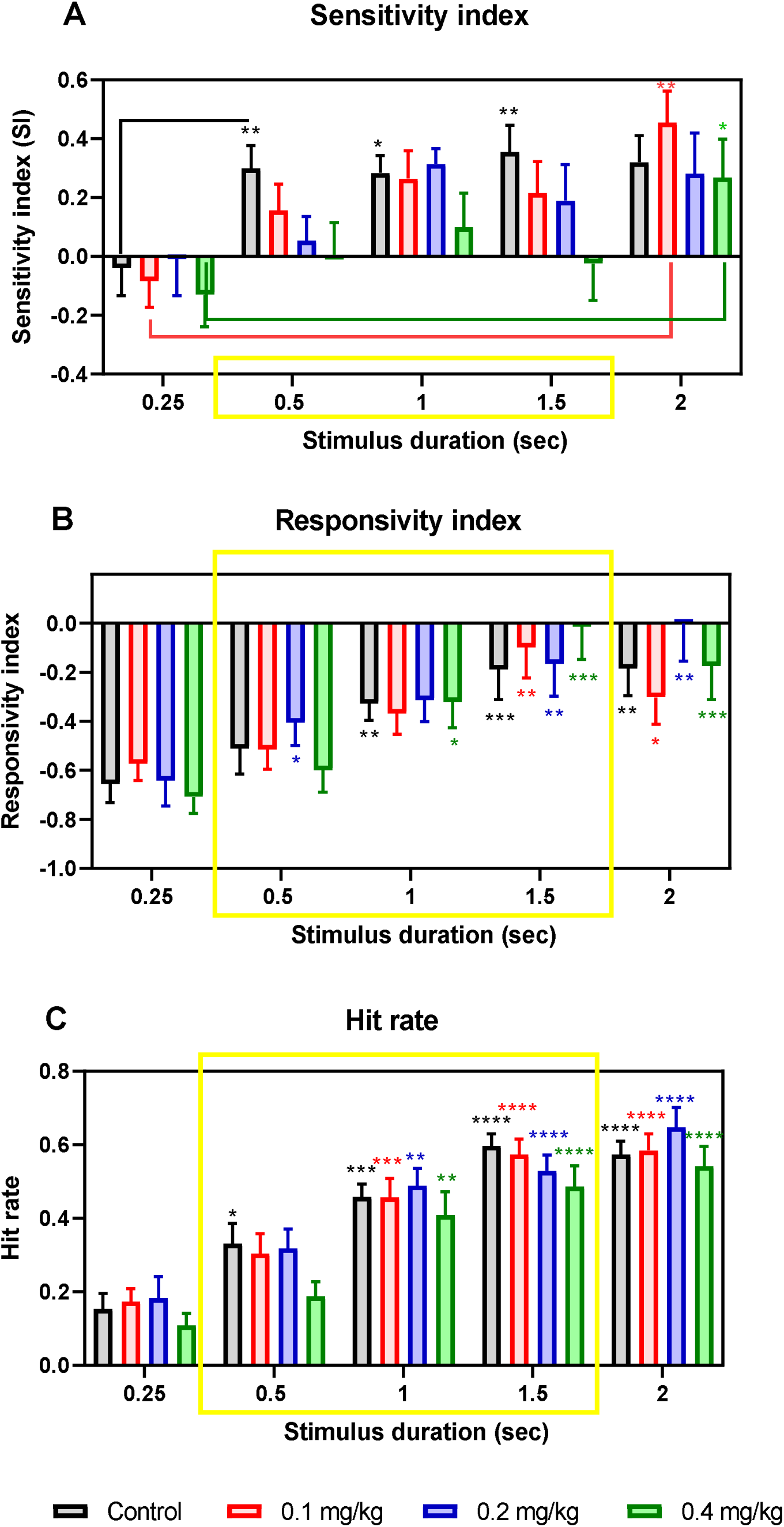
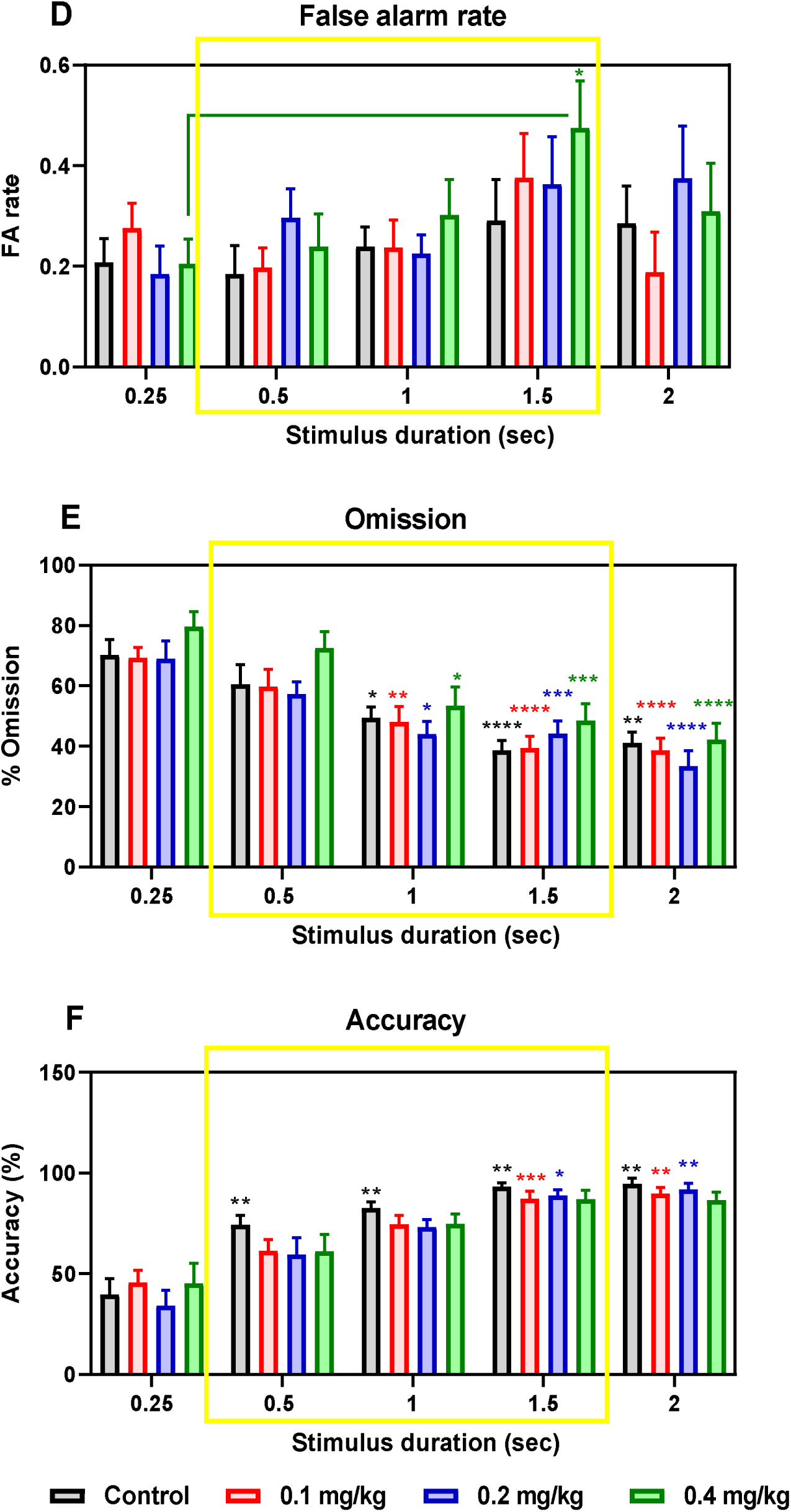
Effects of amphetamine on 5C-CPT performance under each reduced SD (0.25, 0.5, 1, 1.5 and 2s). (A) Sensitivity index. (B) Responsivity index. (C) Hit rate. (D) False alarm rate. (E) Omission. (F) Accuracy (n=14). * *p* < 0.05, ** *p* < 0.01, *** *p* < 0.001 compared to vehicle 0.25s SD (Dunnet). Red asterisks stand for the significance differences between 0.1mg/kg AMPH treated rats at 0.25s SD and 0.1mg/kg AMPH treated rats at longer SDs. Blue asterisks stand for the significance differences between 0.2mg/kg AMPH treated rats at 0.25s SD and 0.2 mg/kg AMPH treated rats at longer SDs. Green asterisks stand for the significance differences between 0.4mg/kg AMPH treated rats at 0.25s SD and 0.4 mg/kg AMPH treated rats at longer SDs. Measures are shown as observed mean ±SEM.

#### Responsivity index and Omission (Fig. 5 B and E)

A two-way repeated measures ANOVA revealed a significant SD effect on responsivity index (F (2.643, 132.8) = 23.90, *p* < 0.0001) and omission (F (3.304, 171.8) = 43.93, *p* < 0.0001) and no significant treatment effect. Dunnett’s multiple comparison *post hoc* test revealed a significant increase in baseline responsivity index and decrease in baseline omission at 1.0s (*p*< 0.01 and *p*< 0.05, respectively), 1.5s (*p*< 0.001 and *p*< 0.0001, respectively) and 2.0s (*p*< 0.01 and *p*< 0.01, respectively) SDs when compared to saline treated rats at 0.25s, indicating that rats were more responsive to target and non-target trials during the longer SDs.

#### Hit rate (Fig. 5C)

A two-way repeated measures ANOVA revealed a significant SD effect (F (3.311, 172.2) = 73.19, *p* < 0.0001) on hit rate and no significant treatment effect. Dunnett’s multiple comparison *post hoc* test revealed a significant increase in baseline hit rate at 0.5s (*p*< 0.05), 1.0s (*p*< 0.001), 1.5s (*p*< 0.0001) and 2.0s (*p*< 0.0001) SDs when compared to baseline (vehicle) hit rate at 0.25s SD, suggesting that hit rate depends on SD length. The highest dose of amphetamine showed a tendency towards reducing hit rate at all the SDs. This was accompanied by a tendency towards increasing false alarm rate at 1.5s SD.

#### False alarm rate (Fig. 5D)

A two-way repeated measures ANOVA revealed a significant SD effect (F (2.599, 130.6) = 3.899, *p* < 0.05) on false alarm rate and no significant treatment effect. Dunnett’s multiple comparison *post hoc* test revealed a significantly increased false alarm rate in rats treated with the highest dose of amphetamine at 1.5s (*p*< 0.05) SDs compared with FA rate of rats treated with the highest dose of amphetamine at 0.25s SD, suggesting that FA rate increases with the length of the SD. Despite this the increased baseline FA rate remained stable at 0.25s and 0.5s SDs and only showed a slight increase at 1.5s SD, remaining stable 1.5s and 2s SD. In other words, reducing the SD reduces HR and, to a lesser extent, FAR (it reduces responsivity to target and non-target trials). The highest dose of amphetamine showed a tendency towards increasing false alarm rate at 1.5s SD, however, it did not reach a significant level.

#### Accuracy (Fig. 5F)

A two-way repeated measures ANOVA revealed a significant SD effect (F (2.504, 25.04) = 32.41, *p* < 0.0001) on accuracy and no significant treatment effect. Dunnett’s multiple comparison *post hoc* test revealed a gradual increase in accuracy as the SDs lengthens (0.5s, *p*< 0.01; 1.0s, *p*< 0.01; 1.5s, *p*< 0.01; 2.0s, *p*< 0.01) when compared to baseline accuracy at 0.25s SD, suggesting that performance accuracy depends on SD. The highest dose of amphetamine also showed a tendency towards increasing omission and decreasing accuracy at 0.5s, 1s and 1.5s SDs, suggesting an impairment in attentional processes.

#### Correlational analysis

Oldham’s method revealed a significant positive association between baseline sensitivity index, hit rate, False alarm rate or omission and the change following administration of the highest dose of amphetamine at 1s SD indicative of rate-dependent effects (Fig.6). The highest dose of AMPH did not change responsivity index at 1s SD in a rate dependent manner, suggesting that the change in sensitivity index at 1s SD was due to change in Hit rate and false alarm rate.

**Figure 6:**
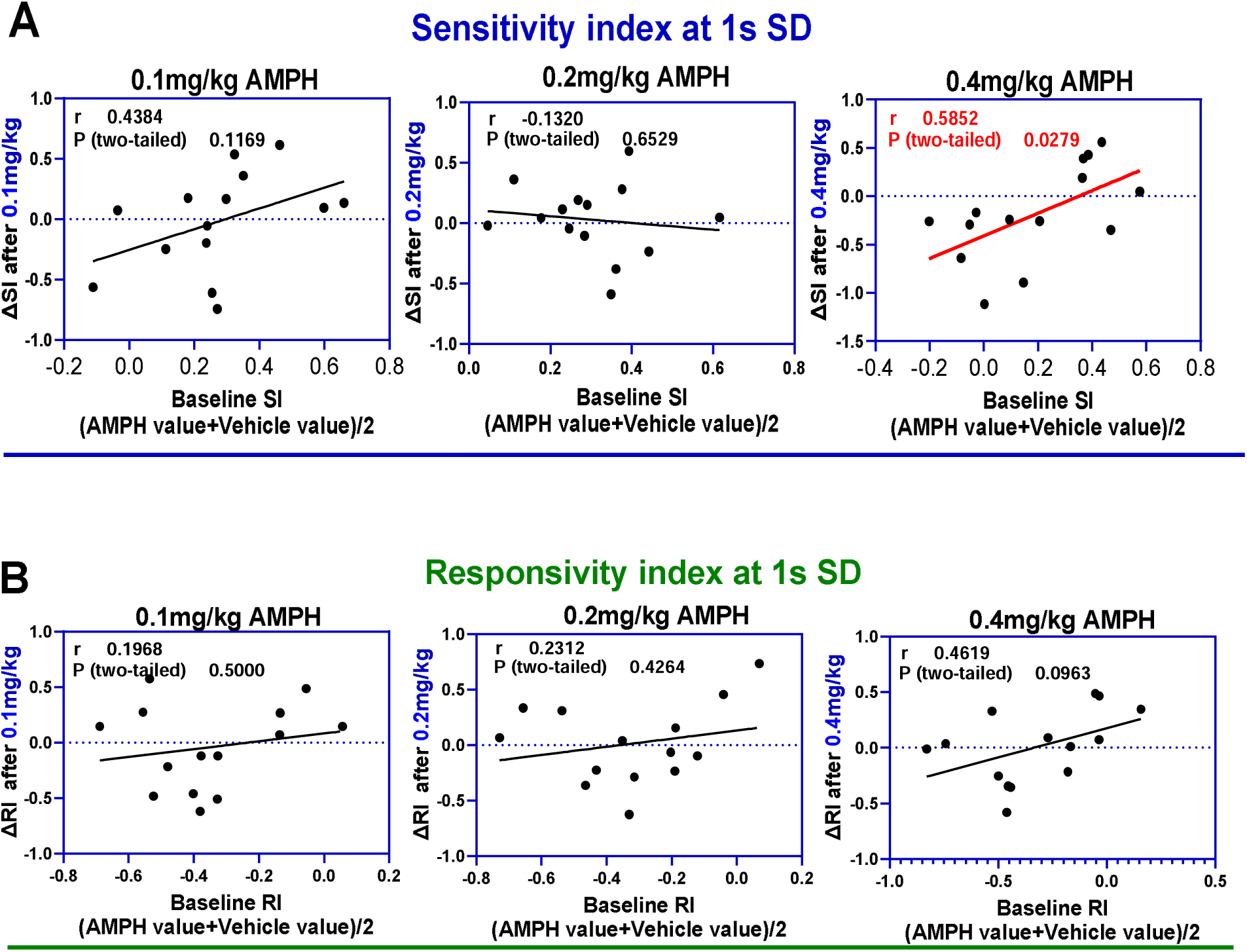
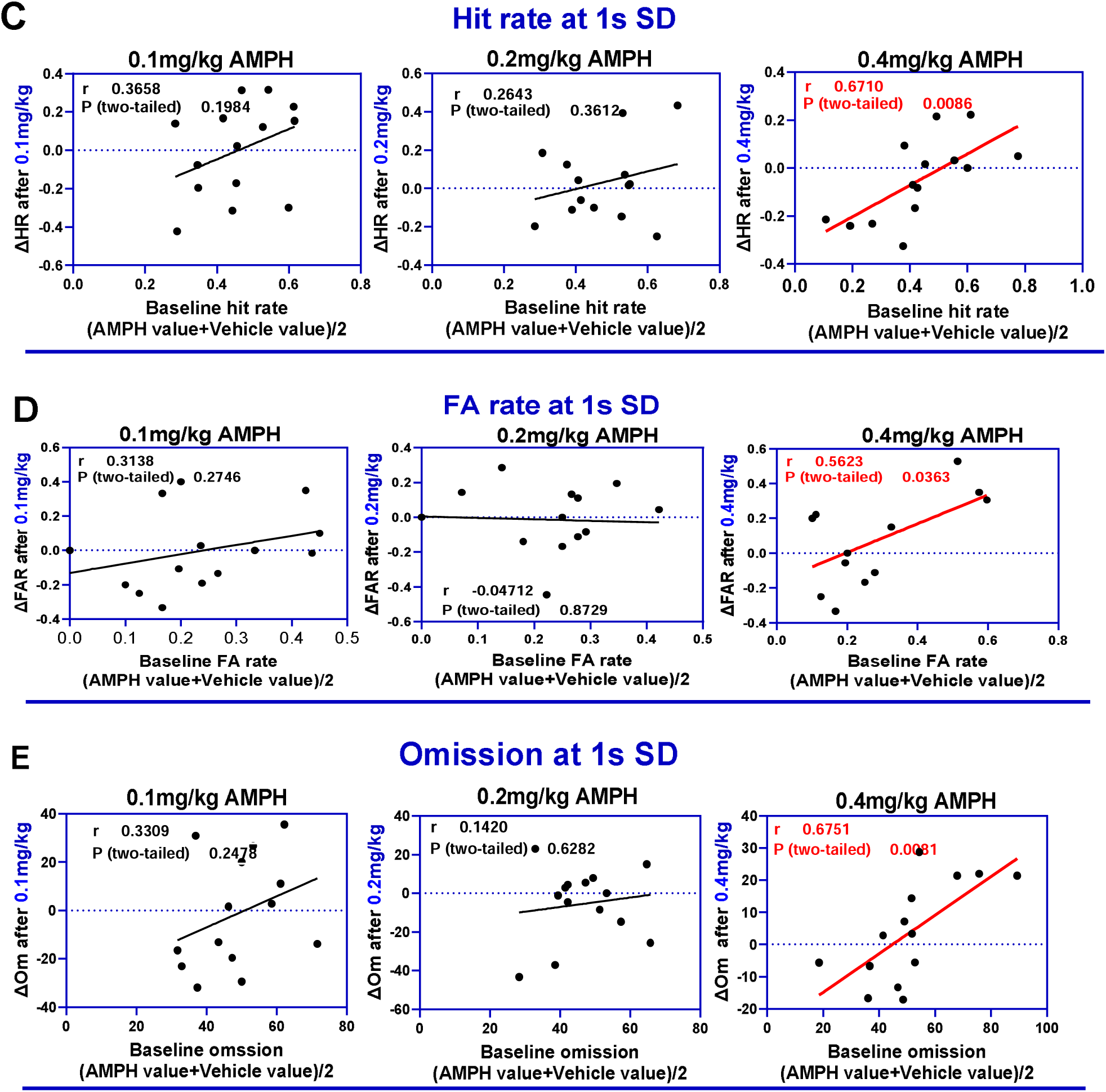
Relationship between baseline 5C-CPT performance measures and the drug induced change at 1s SD using Oldham method (n=14). (A) SI. (B) RI. (C) Hit rate. (D) FA rate. (E) Omission. The red regression lines represent the statistically significant relationships.

Oldham’s method revealed a rate-dependent relationship between baseline accuracy and the change following administration of 0.1 or 0.4 mg/kg AMPH at 1.5s SD (Fig. 7).

**Figure 7:**
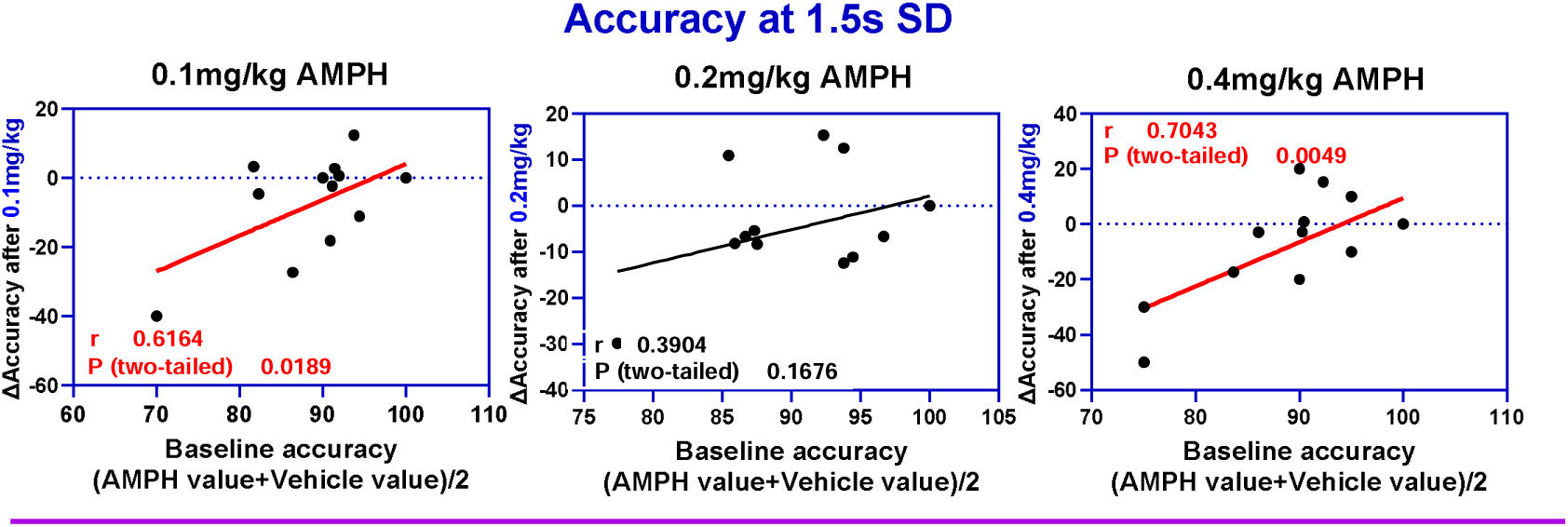
Relationship between baseline accuracy and the drug induced change at 1.5s SD using Oldham method (n=14). The red regression lines represent statistically the significant relationships.

### Correlation between change in 5C-CPT performance measures at each SD following administration of AMPH

Correlational analysis was used to understand the relationship between change in sustained attention and change in responsivity following administration of AMPH at each SD.

**Relationship between change in sustained attention (**ΔSI) **and change in responsivity index** (ΔRI) **following administration of the highest dose of AMPH:** Correlational analysis was used to understand if the ΔSI at 0.4 mg/kg AMPH was due to ΔHR or ΔFAR or both (ΔRI). This was done in three steps: 1) ΔSI was plotted against ΔRI at 0.4mg/kg. 2) ΔSI was plotted against ΔHR at 0.4 mg/kg. 3) ΔSI was plotted against ΔFAR at 0.4 mg/kg. Correlational analysis revealed no association between ΔSI and ΔRI following administration of the highest dose of AMPH (Fig. 8), suggesting that ΔSI was not due to ΔRI. Only two rats showed improvement in SI following administration of AMPH with eights rats showing an impairment in performance. Correlational analysis revealed no association between ΔHR and ΔSI (Fig 8B). Correlational analysis revealed a significant, strong negative relationship between ΔFAR and ΔSI (Fig. 8C). Together, these findings suggest that the ΔSI was mainly due to change in ΔFAR at 0.5s SD.
**Relationship between change in sustained attention and change in responsivity index at 1.0s SD following administration of the highest dose of AMPH:** Here, correlational analysis revealed: 1) no relationship between ΔRI and ΔSI; 2) a significant, moderate positive association between ΔSI and ΔHR and a strong negative association between ΔSI and ΔFAR, suggesting that the ΔSI following administration of 0.4 mg/kg at 1.0s SD was due to ΔHR and ΔFAR.
**Relationship between change in sustained attention and change in responsivity index at 1.5s SD following administration of the highest dose of AMPH:** Here, correlational analysis revealed: 1) a significant strong negative association between ΔSI and ΔRI following administration of the highest dose of AMPH (Fig. 10), suggesting that the ΔSI was due ΔRI. 9 0ut of 14 rats showed impairment in SI; 2) no association between ΔSI and ΔHR; and 3) a significant negative association between ΔSI and ΔFAR.

**Figure 8:**
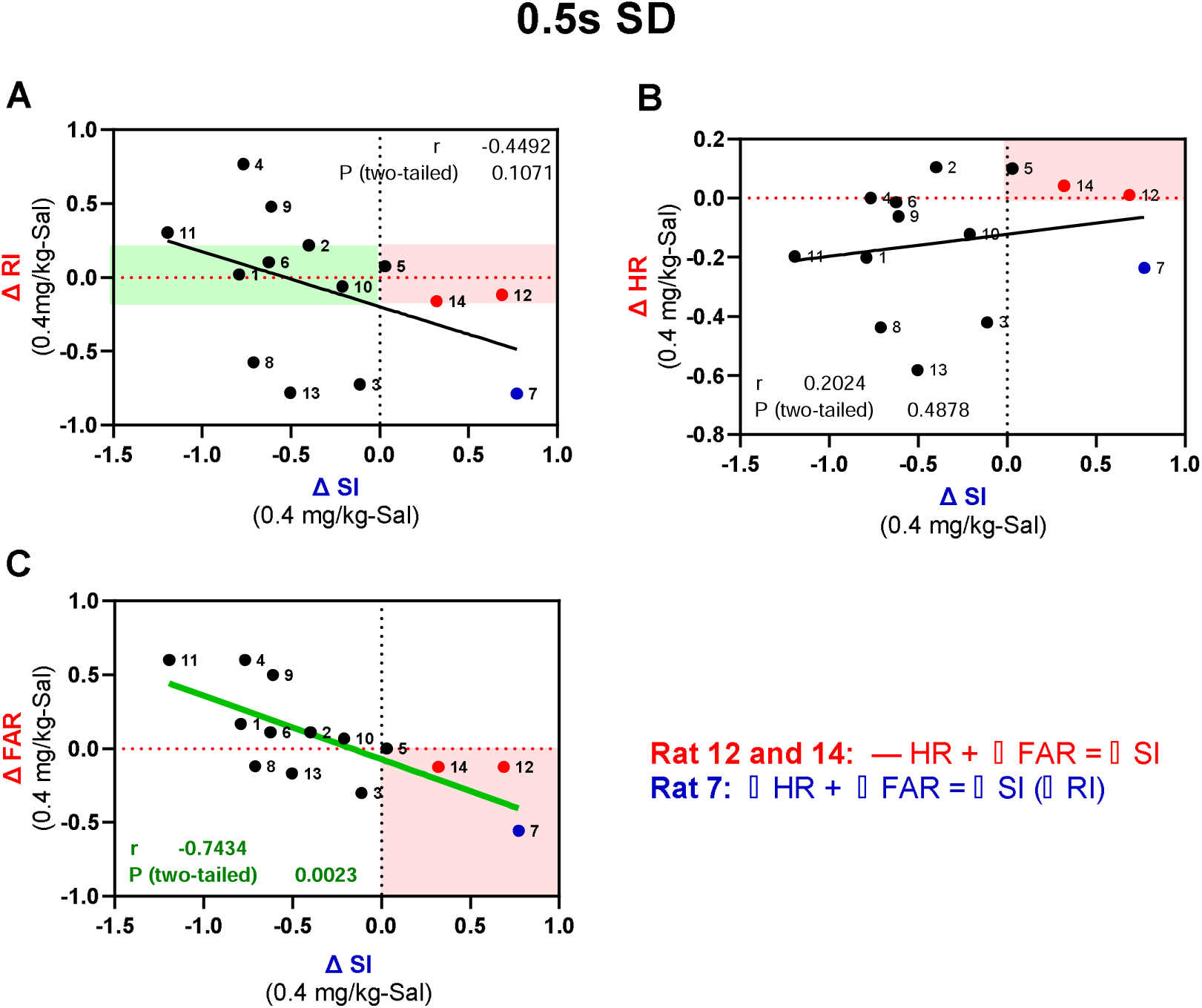
Correlation between 5C-CPT performance measures (n=14). (A) Correlation between ΔSI and ΔRI. (B) Correlation between ΔSI and ΔHR. (C) Correlation between ΔSI and ΔFAR. Task performance measure abbreviations indicate Sensitivity index (SI), Responsivity index (RI), Hit rate (HR), false alarm rate (FAR). Symbols indicate performance improvement (↑), performance impairment (↓) or no effect (□).

**Figure 9:**
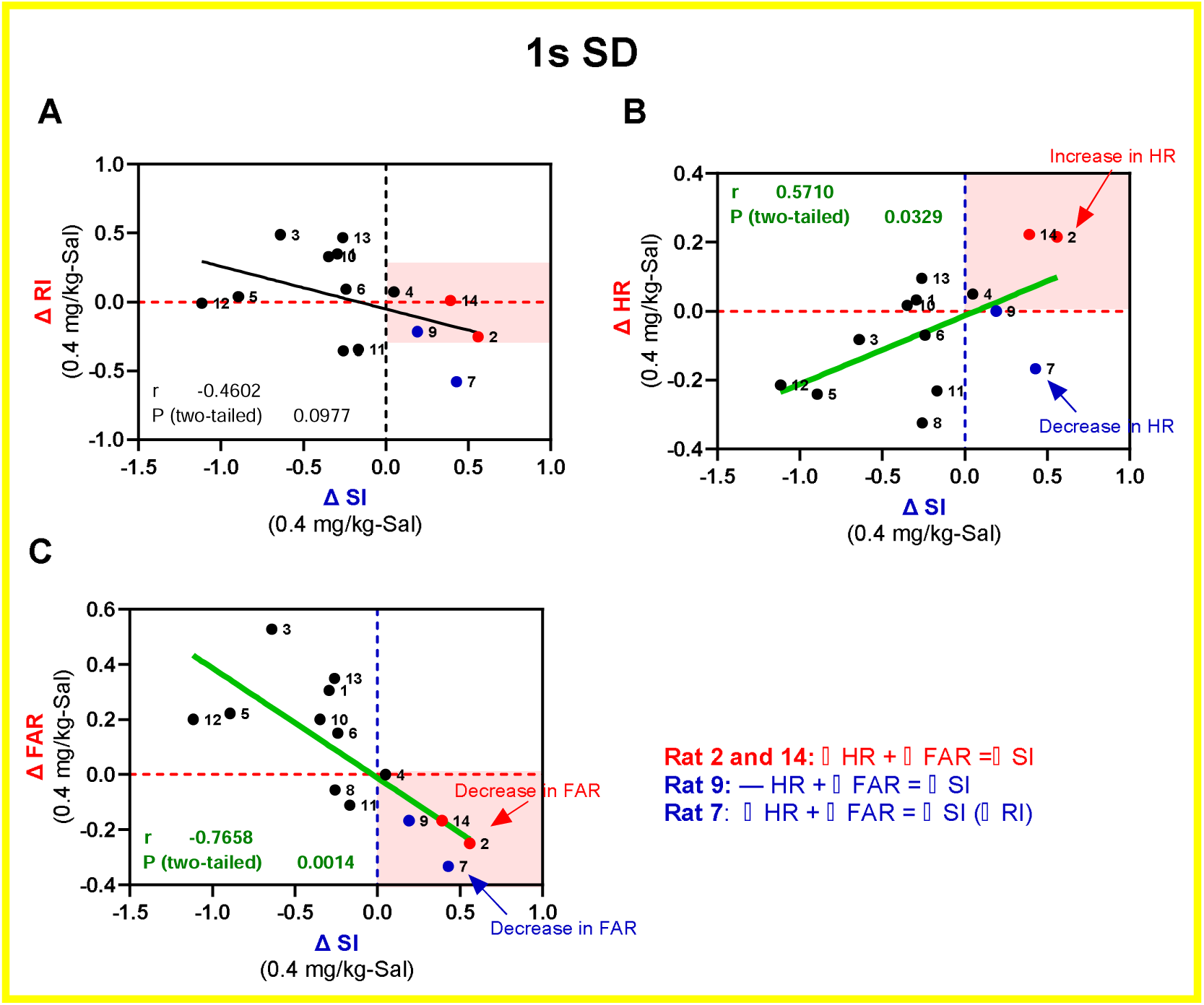
Correlation between change in 5C-CPT performance measures following administration of 0.4 mg/kg AMPH at 1s SD (n=14). (A) Correlation between ΔSI and ΔRI. (B) Correlation between ΔSI and ΔHR. (C) Correlation between ΔSI and ΔFAR. Task performance measure abbreviations indicate Sensitivity index (SI), Responsivity index (RI), Hit rate (HR), false alarm rate (FAR). Symbols indicate performance improvement (↑), performance impairment (↓) or no effect (□).

**Figure 10:**
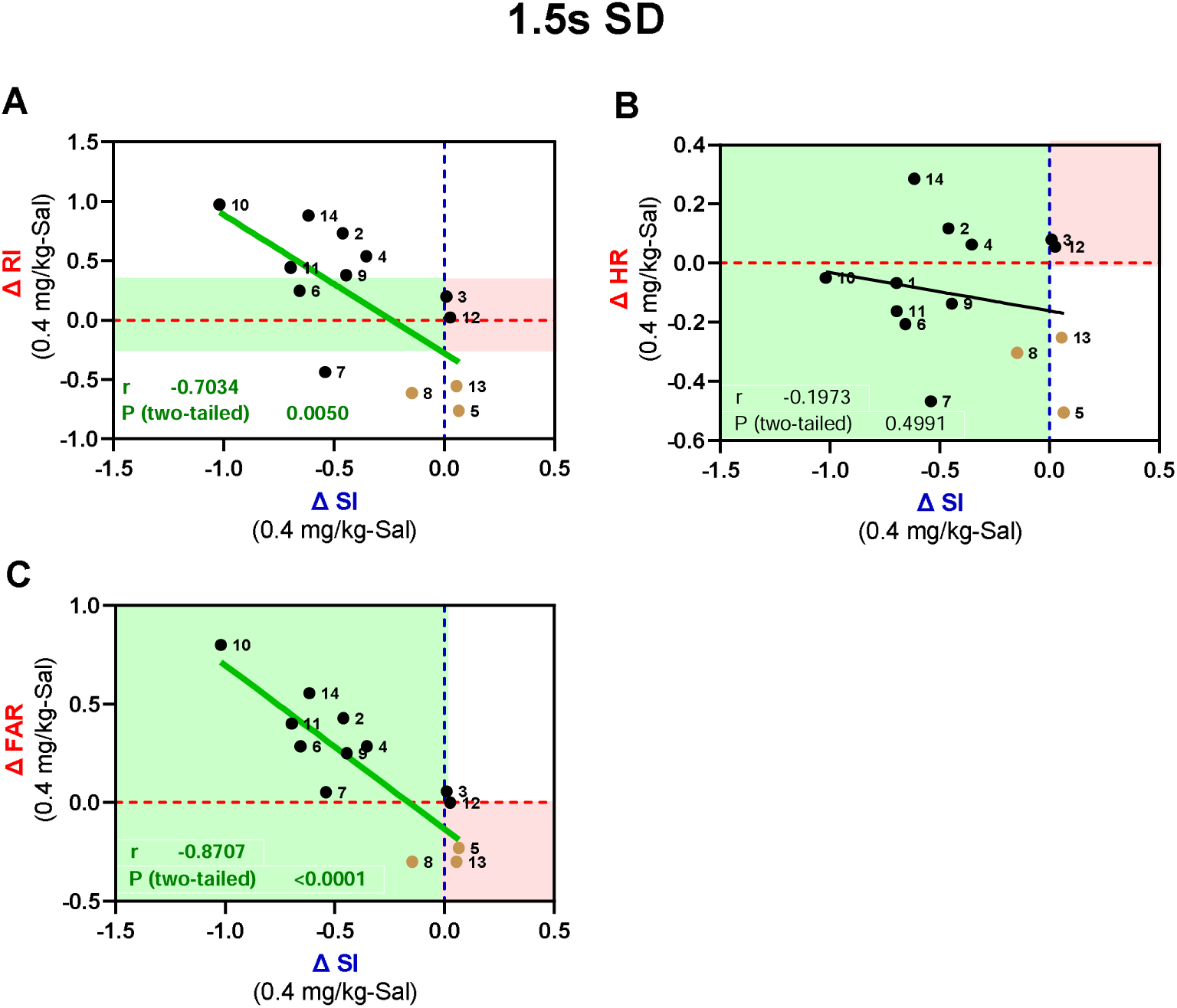
Correlation between change in 5C-CPT performance measures following administration of 0.4 mg/kg AMPH at 1.5s SD (n=14). (A) Correlation between ΔSI and ΔRI. (B) Correlation between ΔSI and ΔHR. (C) Correlation between ΔSI and ΔFAR. Task performance measure abbreviations indicate Sensitivity index (SI), Responsivity index (RI), Hit rate (HR), false alarm rate (FAR). Symbols indicate performance improvement (↑), performance impairment (↓) or no effect (□).

**Figure 11:**
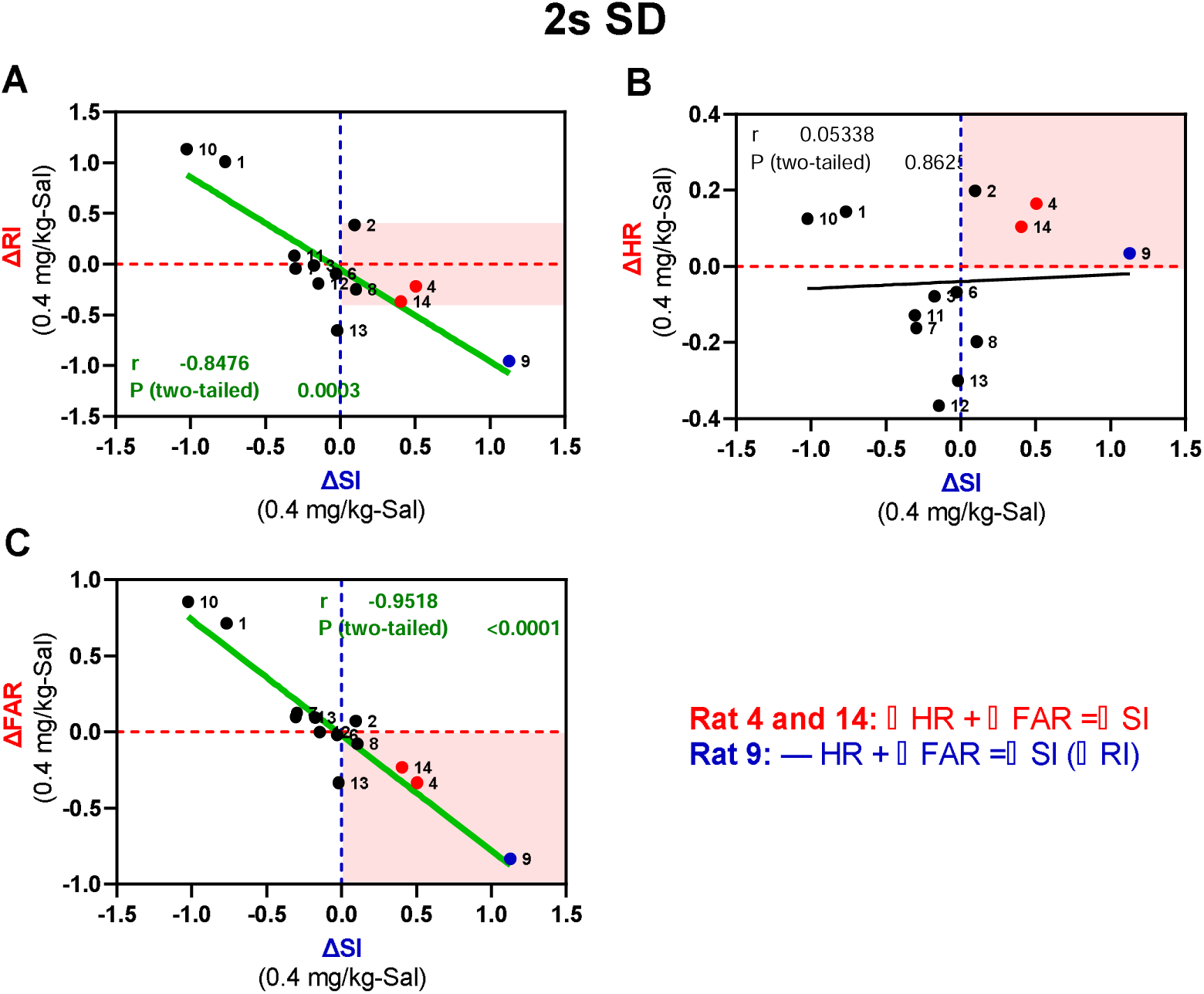
Correlation between change in 5C-CPT performance measures following administration of 0.4 mg/kg at 2s SD (n=14). (A) Correlation between ΔSI and ΔRI. (B) Correlation between ΔSI and ΔHR. (C) Correlation between ΔSI and ΔFAR. Task performance measure abbreviations indicate Sensitivity index (SI), Responsivity index (RI), Hit rate (HR), false alarm rate (FAR). Symbols indicate performance improvement (↑), performance impairment (↓) or no effect (□).

**Figure 12:**
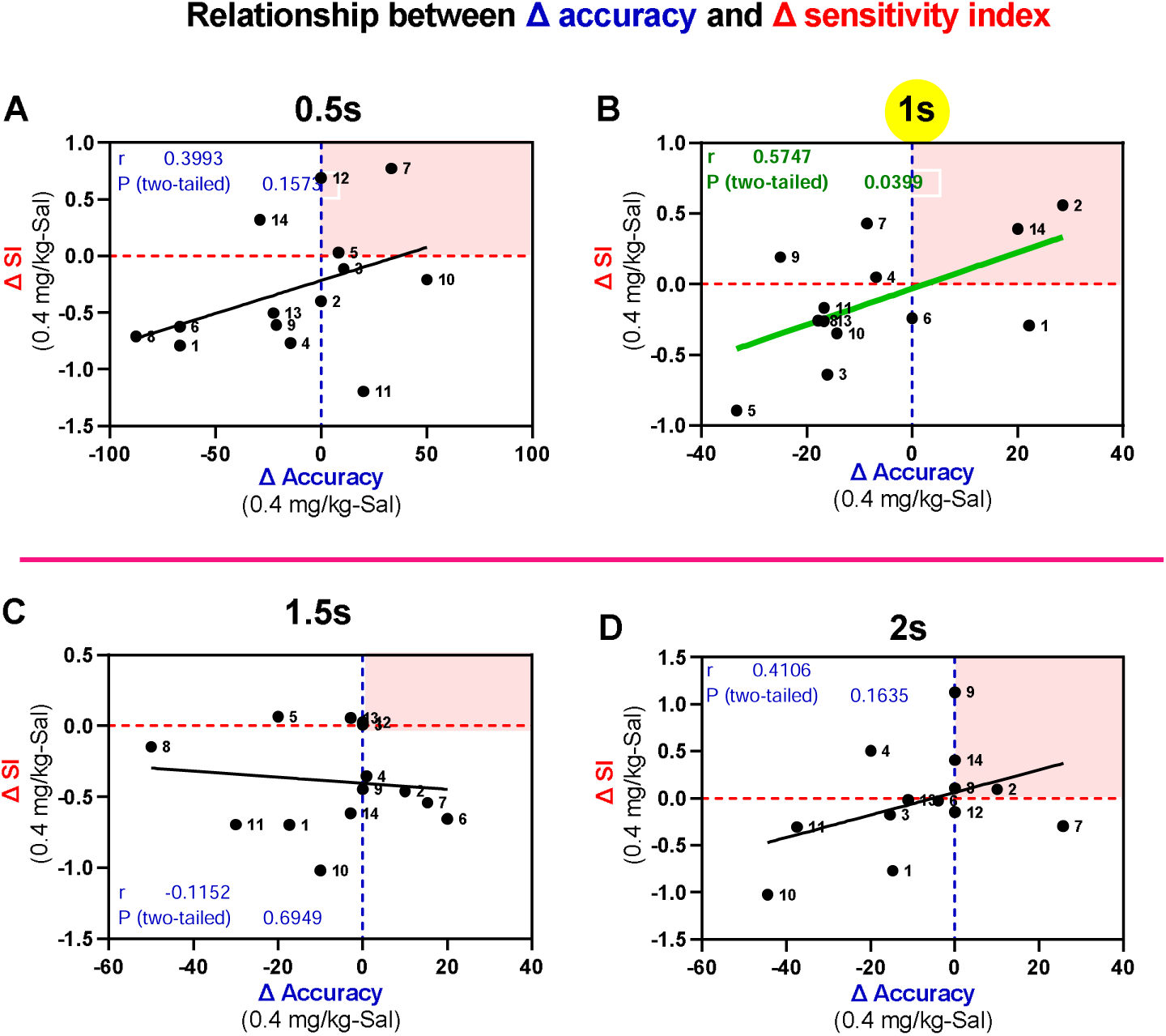
Correlation between Δaccuracy and ΔSI following administration of the highest dose of AMPH (n=14). (A) 0.5s. (B) 1s. (C) 1.5s. (D) 2s.

### Relationship between change in sustained attention and change in responsivity index following administration of the highest dose of AMPH at 2s SD

Correlational analysis revealed a significant, negative association between ΔSI and ΔRI following administration of the highest dose of AMPH. Only two rats showed improvement in performance following administration of the highest dose of AMPH. Correlational analysis also revealed no association between ΔSI and ΔHR and a significant negative association between ΔFAR and ΔSI.

Together, these findings suggest that ΔSI at 0.5 and 1.0s SD following administration of 0.4 mg/kg AMPH was not due to ΔRI, however, ΔSI at longer SDs (1.5s and 2.0s SDs) following administration of 0.4 mg/kg AMPH was due to ΔRI.

### Relationship between Δaccuracy and ΔSI following administration of the highest dose of AMPH under each reduced SDs

Correlational analysis revealed a significant relationship between Δaccuracy and ΔSI at 1s SD.

Relationship between Δaccuracy and ΔRI following administration of the highest dose of AMPH at each SD is shown in Fig. 13. Correlational analysis revealed no significant relationship between Δaccuracy and ΔRI.

**Figure 13:**
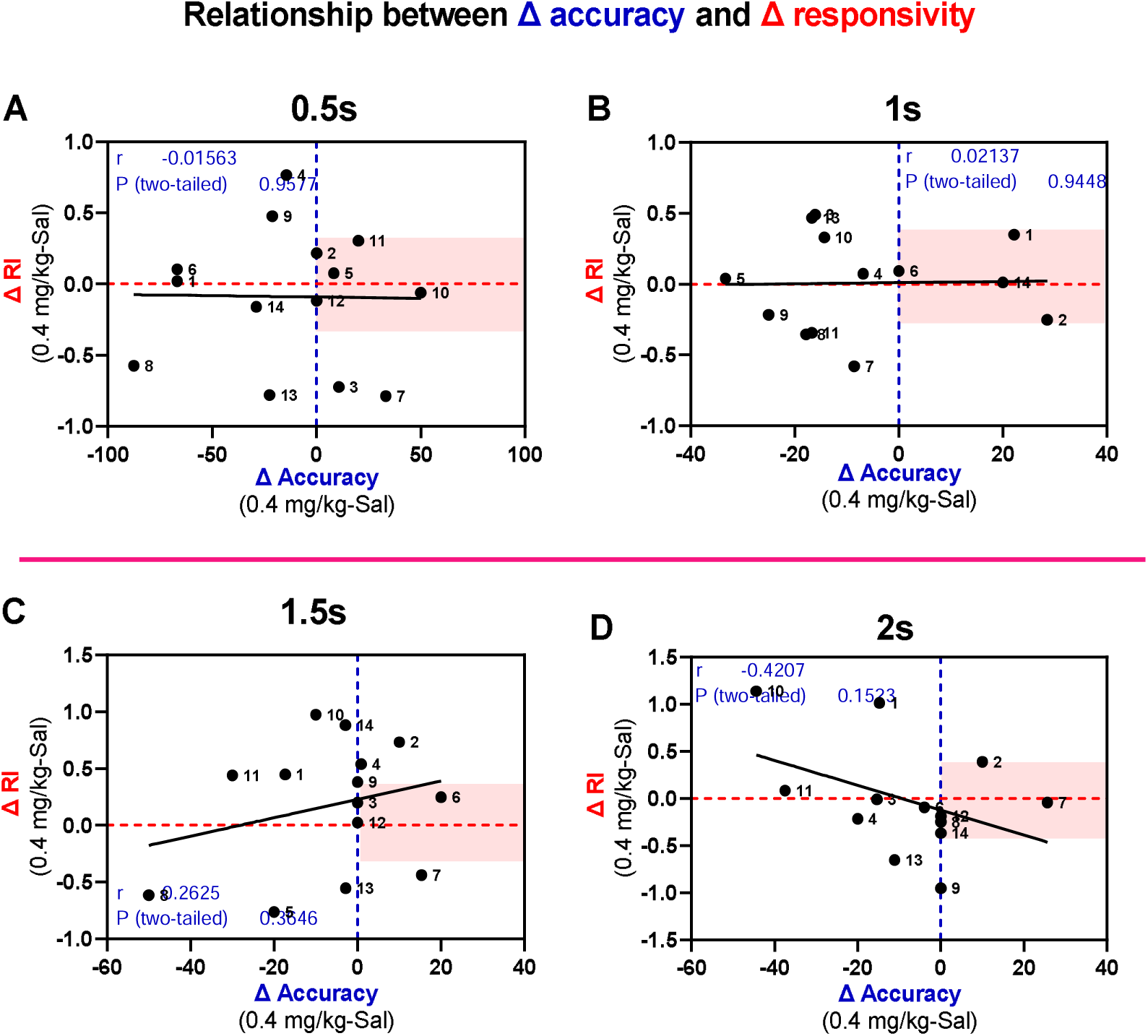
Correlation between Δaccuracy and ΔRI following administration of the highest dose of AMPH at each SD (n=14). (A) 0.5s. (B) 1s. (C) 1.5s. (D) 2s.

The relationship between Δaccuracy and Δomission following administration of the highest dose of AMPH at each SD is shown in Fig. 14. Correlational analysis revealed no significant relationship between Δaccuracy and Δomission. Only two rats showed improvement in accuracy and omission following administration of 0.4 mg/kg AMPH at 1.0s SD and only one rat showed improvement in accuracy and omission at 1.5s and 2.0s SDs.

**Figure 14:**
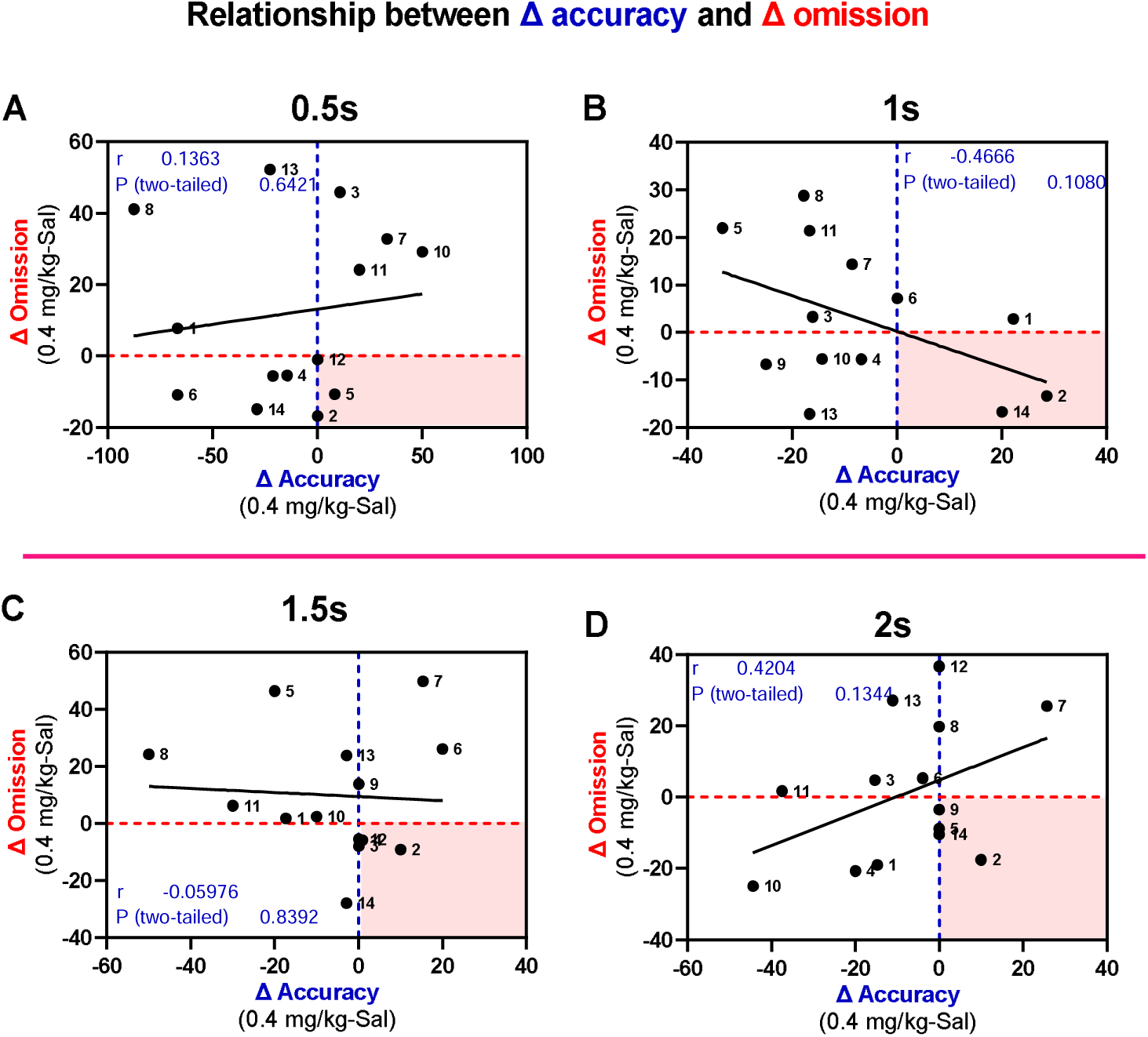
Correlation between Δaccuracy and Δ omission following administration of the highest dose of AMPH under each SD (n=14). (A) 0.5s. (B) 1s. (C) 1.5s. (D) 2s.

### Within-session performance

The within-session performance is shown in Fig. 15, where each session consisted of 4 blocks of 30 trials. The fourth trial block was excluded from the analysis because some of the rats did not complete it.

**Figure 15:**
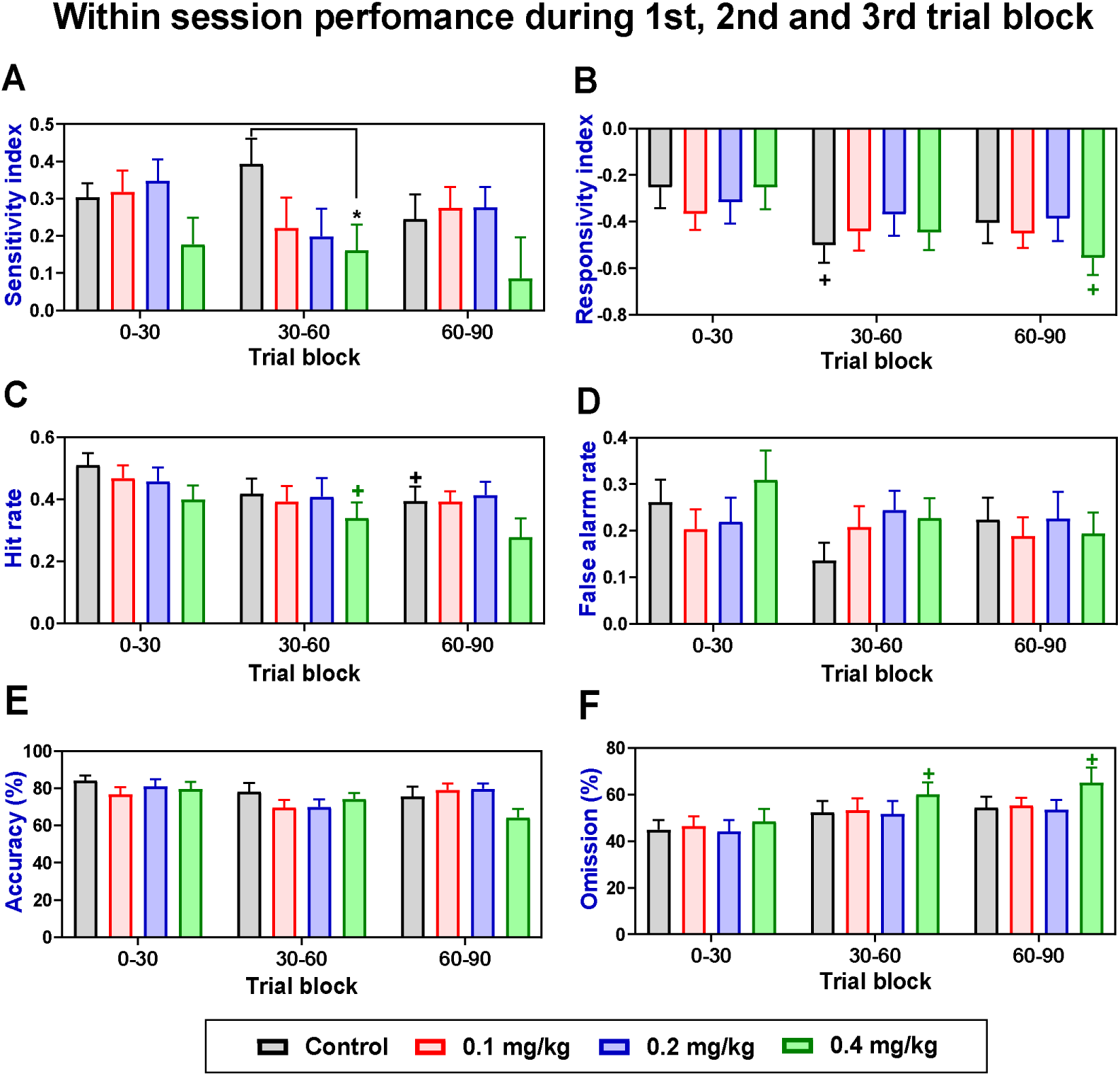
AMPH induced changes on 5C-CPT performance measures over session trials. Data are presented in 30 trial blocks (n=14). Measures are shown as observed mean ±SEM. * *p* < 0.05 compared to control of the same block (Dunnett). +*p* < 0.05 compared to vehicle-treated rats (control) or AMPH treated rats of 1st trial block. A significant difference between vehicle treated rats in the second or third trial block and vehicle treated rats in the first trial block is indicated with a black plus (+). A significant difference between 0.4 mg/kg AMPH treated rats in the second or third trial block and 0.4 mg/kg AMPH treated in the first trial block is indicated with a green plus (+).

#### Sensitivity index (SI)

The within-session performance is shown in Fig.15A, where each session consisted of 4 blocks of 30 trials. A two-way, repeated measures ANOVA revealed no significant effect of trial [F (1.569, 20.40) = 0.9280, *p* = 0.38], a significant treatment effect [F (2.693, 35.01) = 3.158, *p* < 0.05] on sensitivity index but no interaction. Dunnett tests also revealed that the highest dose of AMPH reduced SI (*p*<0.05) in the second trial block compared to control.

#### Responsivity index (RI)

A two-way, repeated measures ANOVA revealed a significant effect of trial [F (1.982, 25.77) = 10.04, *p* < 0.001; Fig. 15B], no treatment effect [F (2.777, 36.10) = 0.4275, *p* = 0.7] and no interaction for responsivity index. Dunnett tests revealed a significant decrease in baseline RI as the session progressed. Baseline RI was significantly lower in the second compared to the first trial block (*p*<0.05). In addition, Dunnett’s revealed a significant decrease in RI of 0.4 mg/kg treated rats in the third compared to the first trial block (*p*<0.05).

#### Hit rate (HR)

A two-way, repeated measures ANOVA revealed a significant effect of trial [F (1.471, 19.13) = 5.832, *p* < 0.05; Fig. 15C] but no treatment effect [F (2.753, 35.79) = 2.307, *p* = 0.09] on hit rate. Dunnett tests revealed a significant decrease in baseline hit rate as the session progressed. Baseline hit rate was significantly lower in the third trial block compared to the first (*p*<0.05). In addition, the Dunnett test revealed a significant decrease in RI of 0.4 mg/kg treated rats in the second trial block compared to the first (*p*<0.05).

#### False alarm rate (FAR)

A two-way, repeated measures ANOVA revealed no significant effect of trial [F (1.851, 24.06) = 2.157, *p* =0.14; Fig. 15D] or treatment [F (2.389, 31.06) = 0.6317, *p* = 0.5] on FAR.

#### Accuracy

A two-way, repeated measures ANOVA revealed no significant effect of trial [F (1.719, 22.35) = 3.011, *p* =0.07; Fig. 15E] or treatment [F (2.298, 29.88) = 1.185, *p* = 0.32] on accuracy.

#### Omission

A two-way, repeated measures ANOVA revealed a significant effect of trial [F (1.706, 22.18) = 7.929, *p* < 0.01; Fig.15F] but not treatment [F (2.796, 36.35) = 1.502, *p* = 0.23] on omission. Dunnett tests revealed that omission in 0.4mg/kg treated rats was significantly higher in the second compared to the first trial block (*p* < 0.05). In addition, omission in 0.4mg/kg treated rats was significantly higher in the third trial block compared with the first (*p* < 0.05) Some rats did not complete 120 trials; therefore, the last trial block was excluded from the analysis.

Correlational analysis using Oldham’s method revealed a significant moderate positive relationship between baseline SI and the change following administration of the highest dose of AMPH, indicative of a rate-dependent effect (Fig. 16A). AMPH did not change other performance measures in a rate-dependent manner (HR, RI, accuracy and omission) (see Supplement 4 Table 2 for an outline of Relationship between baseline 5C-CPT performance measures and AMPH induced change during first, second or third trial block).

**Figure 16:**
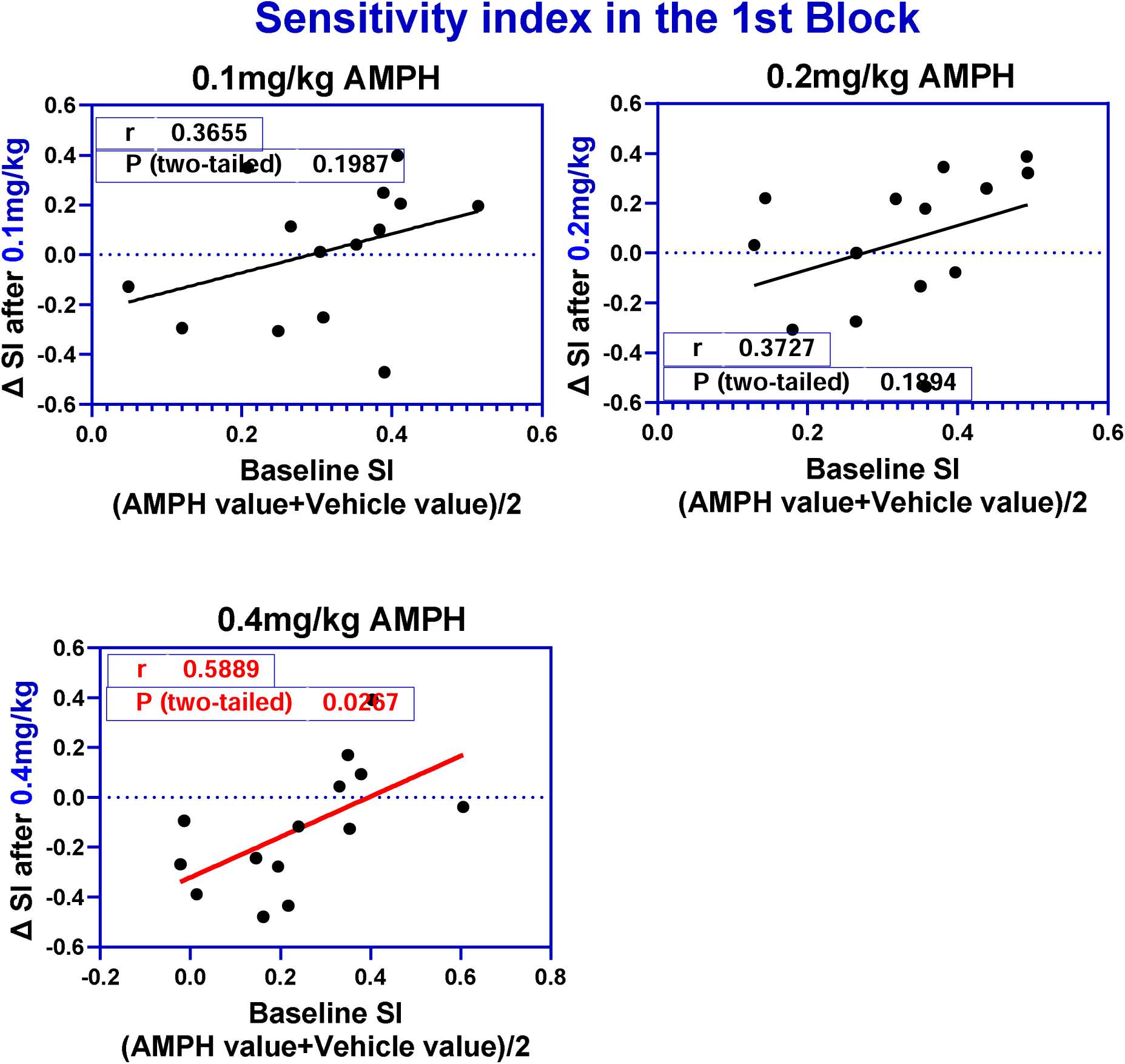
Relationship between baseline sensitivity index and AMPH induced change during the first trial block using Oldham method (n=14). The red regression lines represent the significant relationship.

During the second trial block, Oldham’s method revealed a significant negative relationship between baseline accuracy and the change following administration of 0.4 mg/kg AMPH, indicative of a rate dependent effect (Fig 17).

**Figure 17:**
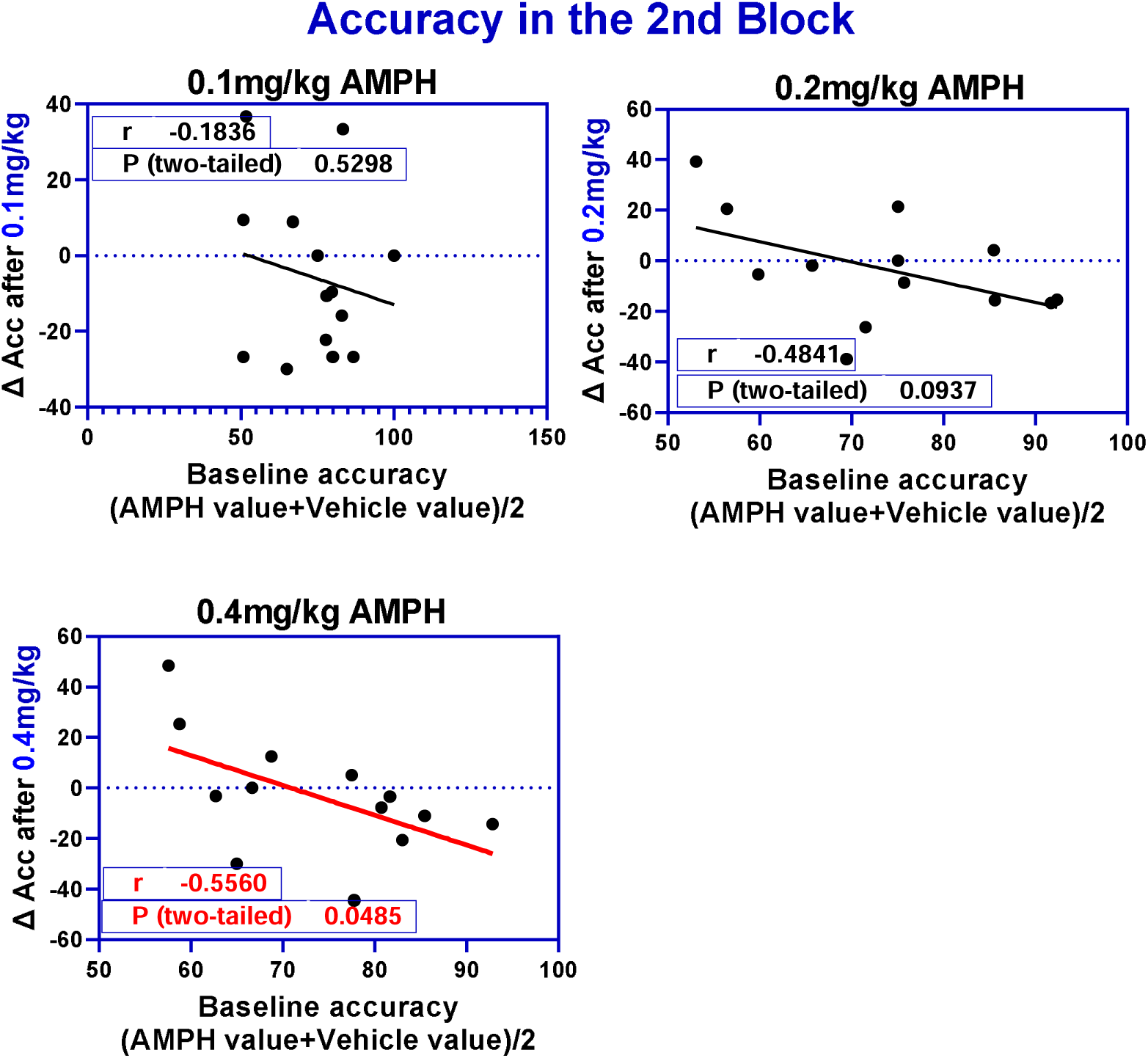
Relationship between baseline accuracy and the drug induced change during the 2^nd^ trial block (n=14). The red regression lines represent the significant relationship.

During the third trial block, 0.2 mg/kg AMPH changed accuracy in a rate-dependent manner. Oldham’s method revealed an inverse relationship between baseline accuracy and magnitude of change following administration of 0.2 mg/kg AMPH (Fig. 18).

**Figure 18:**
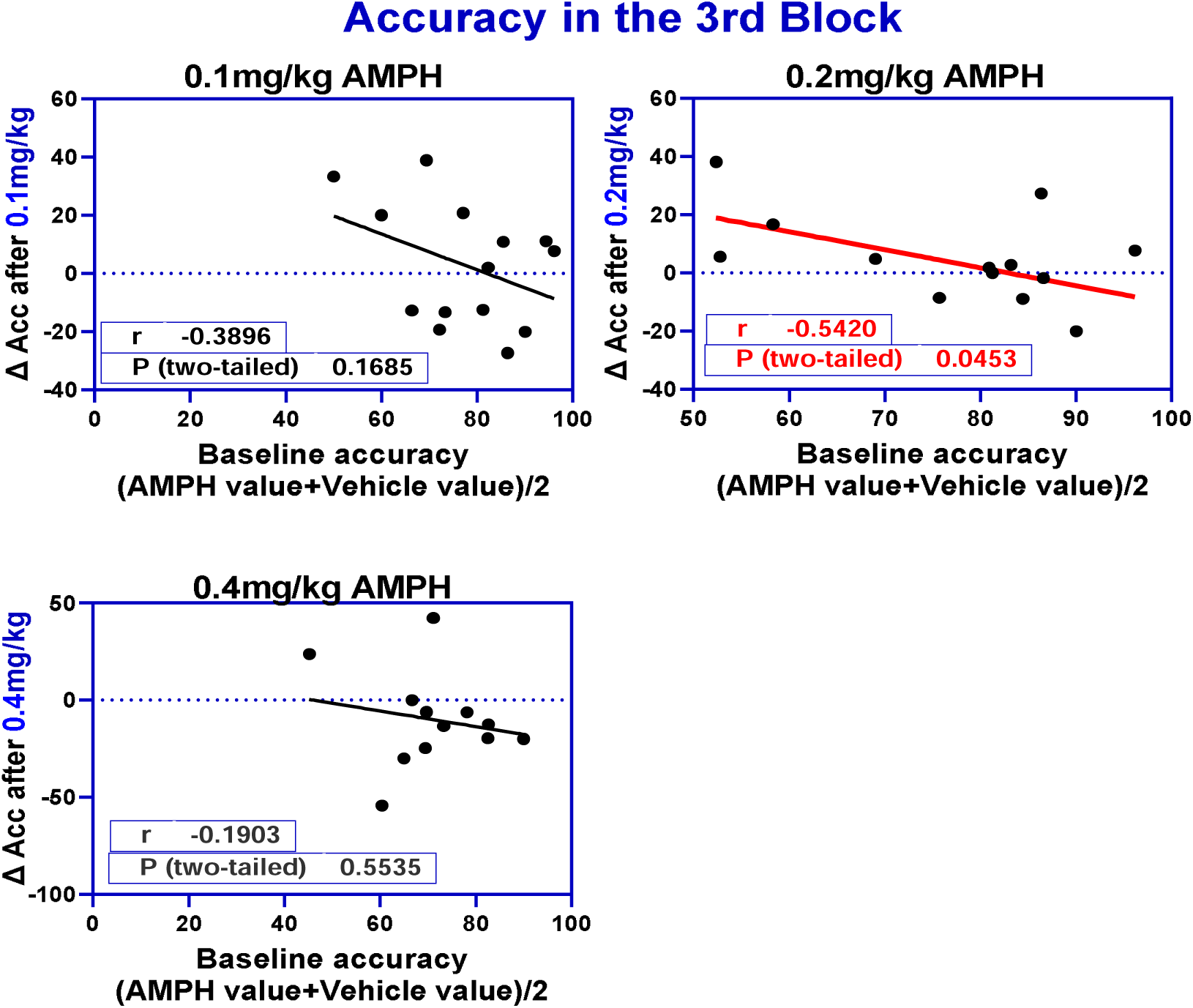
Relationship between baseline accuracy and AMPH induced change during the 3rd trial block using Oldham method (n=14). The red regression lines represent the significant relationship.

### Relationship between Δ SI and Δ RI following administration of 0.4 mg/kg AMPH at each trial block

Correlational analysis revealed no association between change in SI and change in responsivity index at 1^st^, second and third trial block, suggesting that the change in SI was not due to a change in responsivity index (Fig. 19).

**Figure 19:**
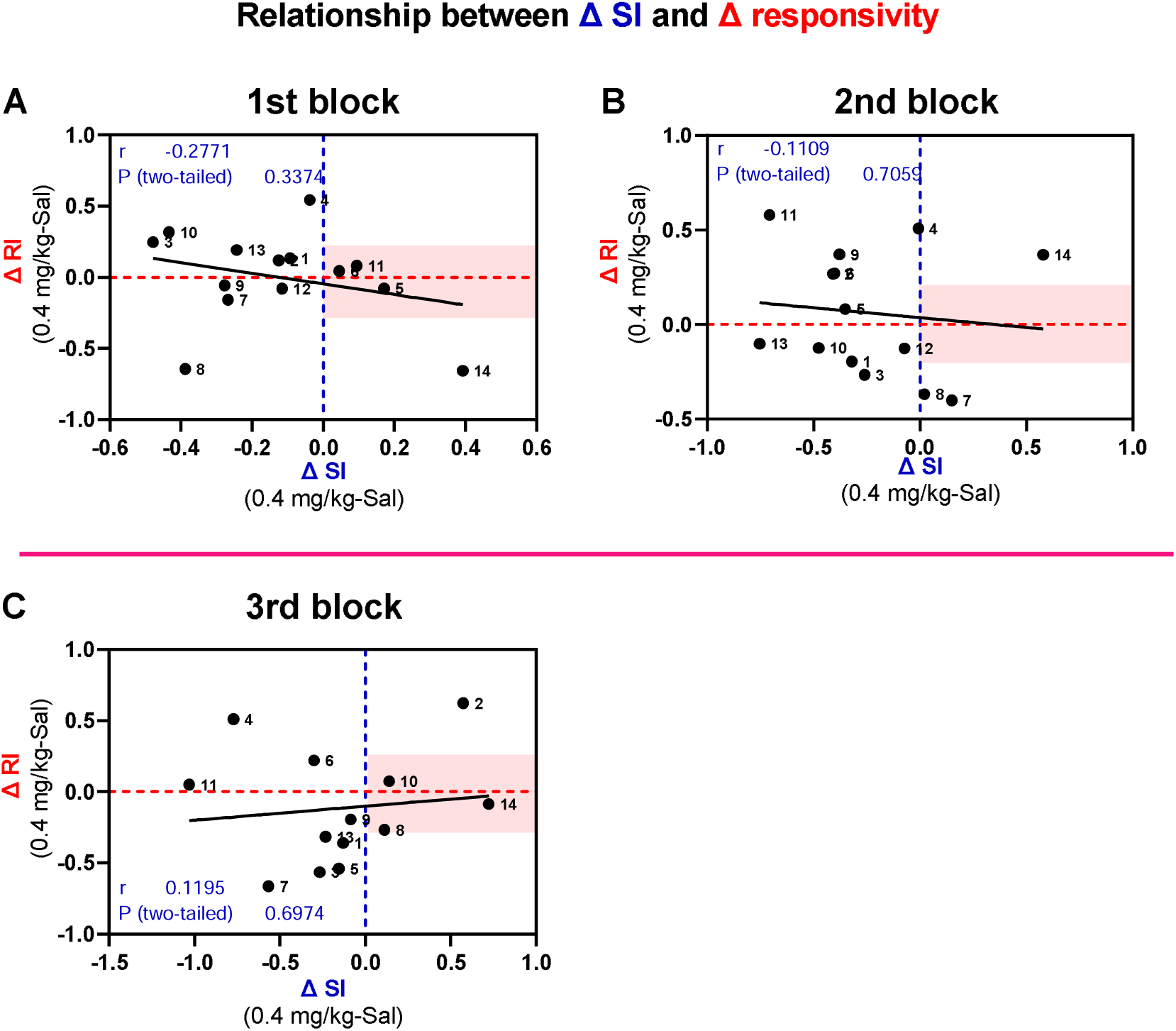
Correlation between change in SI and change in RI following administration of 0.4 mg/kg AMPH (n=14). Task performance measure abbreviations indicate Sensitivity index (SI), Responsivity index (RI). The pink band represents improvement in SI accompanied by small change in RI.

Correlational analysis revealed a significant positive relationship between change in SI and change in HR in the third block, suggesting that the change in SI was due to change in HR (Fig. 20).

**Figure 20:**
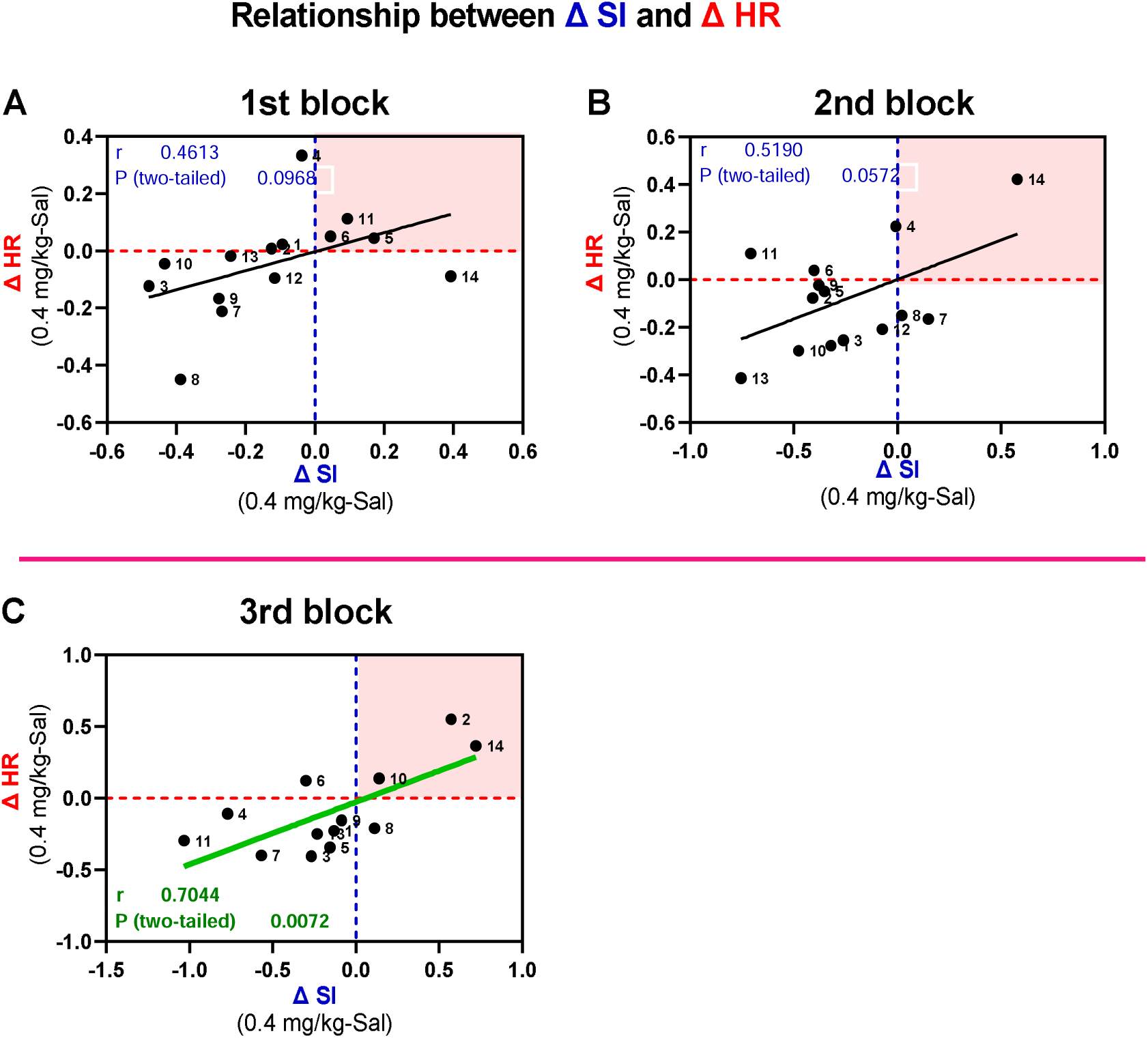
Correlation between change in SI and change in HR following administration of 0.4 mg/kg AMPH (n=14). Task performance measure abbreviations indicate Sensitivity index (SI), Hit rate. The pink square represents improvement in SI due to improvement in HR.

Correlational analysis revealed a significant, negative relationship between change in SI and change in FAR at 1^st^ and 2^nd^ block, suggesting that the change in SI was due to change in FAR (Fig. 21).

**Figure 21:**
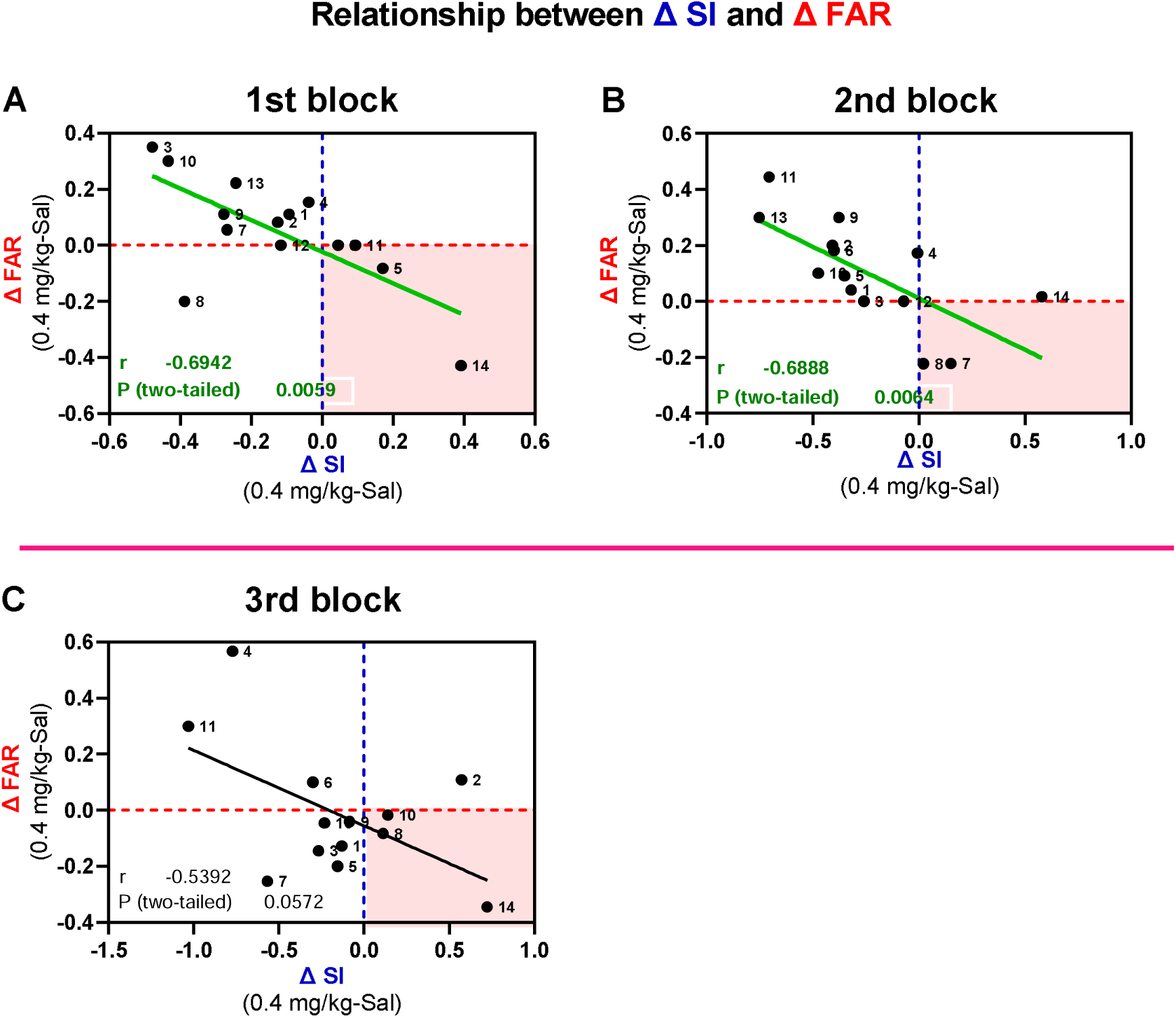
Correlation between change in SI and change in FAR following administration of 0.4 mg/kg AMPH (n=14). Task performance measure abbreviations indicate Sensitivity index (SI), false alarm rate (FAR). The pink square represents improvement in SI due to improvement in FAR.

**Figure 22:**
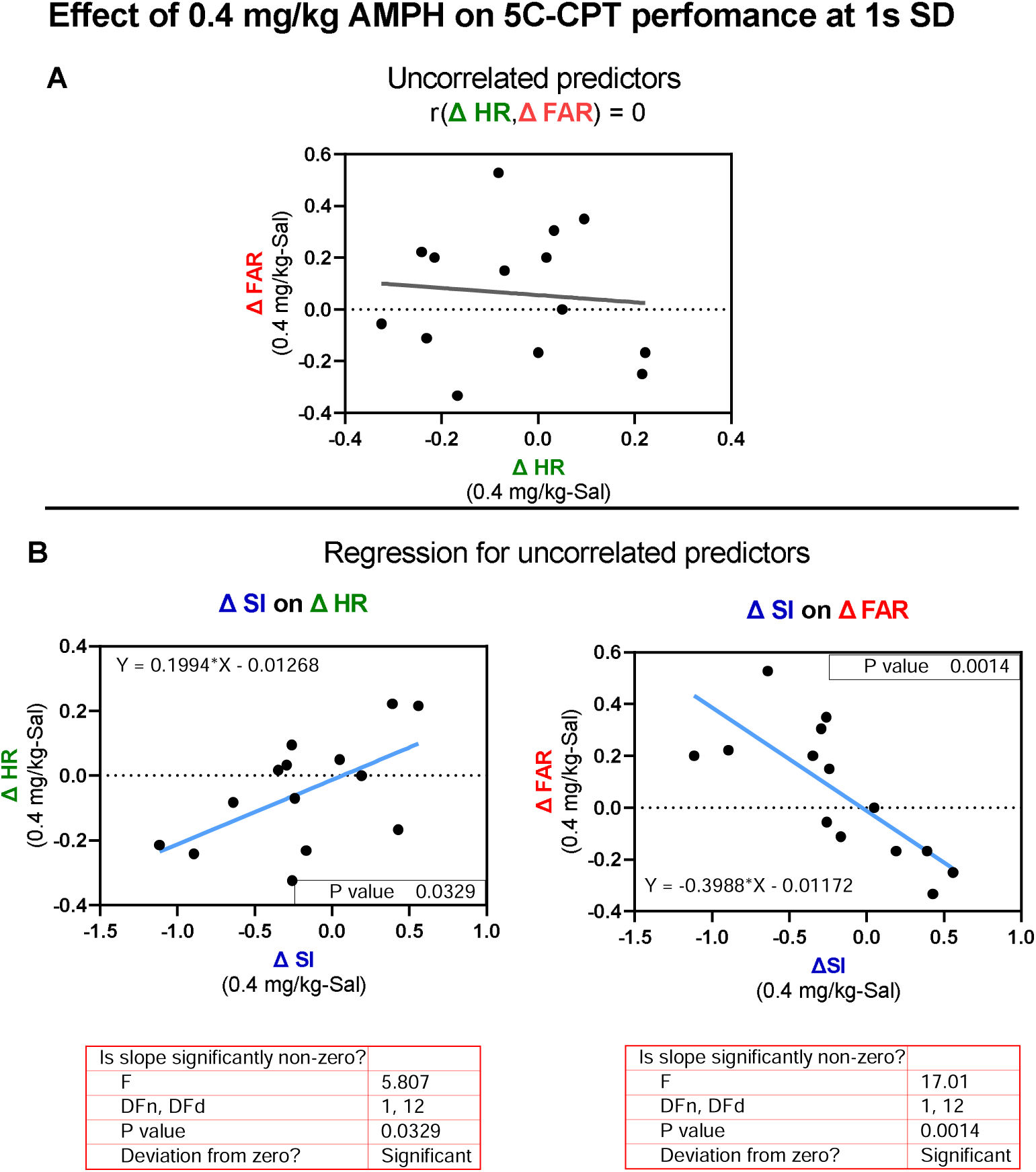
The results of MLR depends on the correlation between ΔHR and ΔFAR. (A) ΔHR and ΔFAR were not correlated with each other (n=14). (B) ΔHR or Δ FAR was correlated with ΔSI.

**Figure 23:**
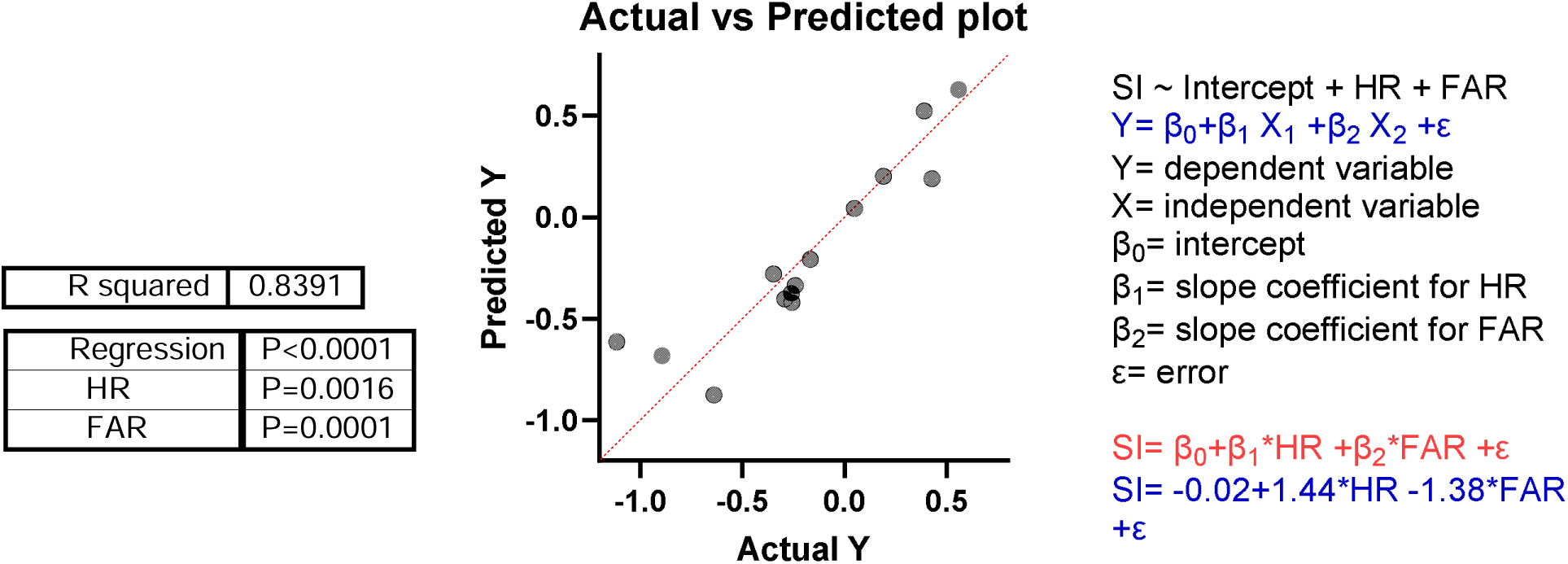
Actual vs predicted plot. R^2^=0.83 suggesting that 83% of the variation in the ΔSI can be explained by the variation in the ΔHR and ΔFAR.

### Multiple linear regression at 1s SD

The independent variables (the change in HR and the change in FAR) are correlated with the dependent variable (SI) and they are not highly correlated with each other, therefore, a multiple linear regression analysis can be used to understand the relationship between these variables.

MLR analysis was conducted to examine the relationship between the independent variables (ΔHR and ΔFAR) and ΔSI following administration of the highest dose of AMPH at 1sSD. Overall, results indicate that the model significantly predicts ΔSI (R2 = 0.83, F (2, 11) = 28.67, p < 0.0001). This model accounts for 83% of the variance observed in ΔSI. A summary of coefficients is presented in Table 6.4.

**Table 6.4.**
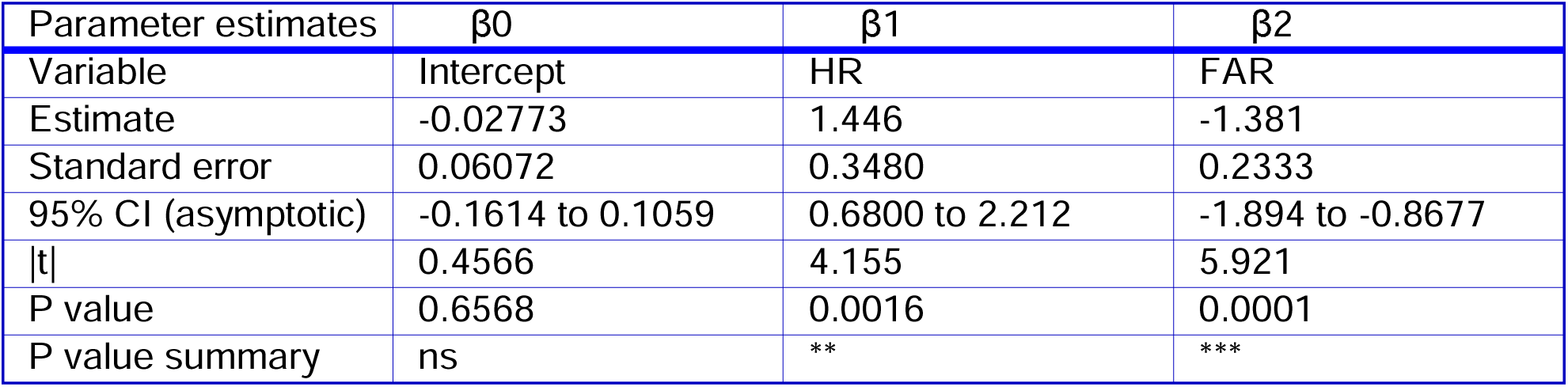
MLR analysis predicting SI. A summary of coefficients.

## Measurements of Oldham’s angles for accuracy

Angle measurements are shown in Fig 24. Amount of space between the regression lines were measured using a protractor. Three angles were measured: ∠1 (blue), ∠2 (green) and ∠3 (red) as shown in Fig 24. Oldham’s angle measurements for the relationship between baseline accuracy and AMPH induced change in the second block are shown in Fig 24A. ∠1 is the space between the highest and the lowest dose (0.4 mg/kg and 0.1 mg/kg regression lines). ∠2 is the space between the highest and the intermediate dose (0.4 mg/kg and 0.2 mg/kg regression lines). ∠3 is the space between 0.2 mg/kg and 0.1 mg/kg lines (Fig 24). The clockwise movement (⟳) of the regression line as the dose increases shows that the highest dose is more effective than the lowest or the intermediate dose at increasing accuracy of low attentive rats and decreasing accuracy of high attentive rats (Fig 24A). Oldham’s angle measurements for the relationship between baseline accuracy and AMPH induced change at the third block are shown in Fig 24B. If two angles are known, then the third angle can be calculated. The angle measurement and the direction of movement (clockwise or counterclockwise) shows how effective is one dose at increasing accuracy of low attentive rats compared with other doses.

**Figure 24:**
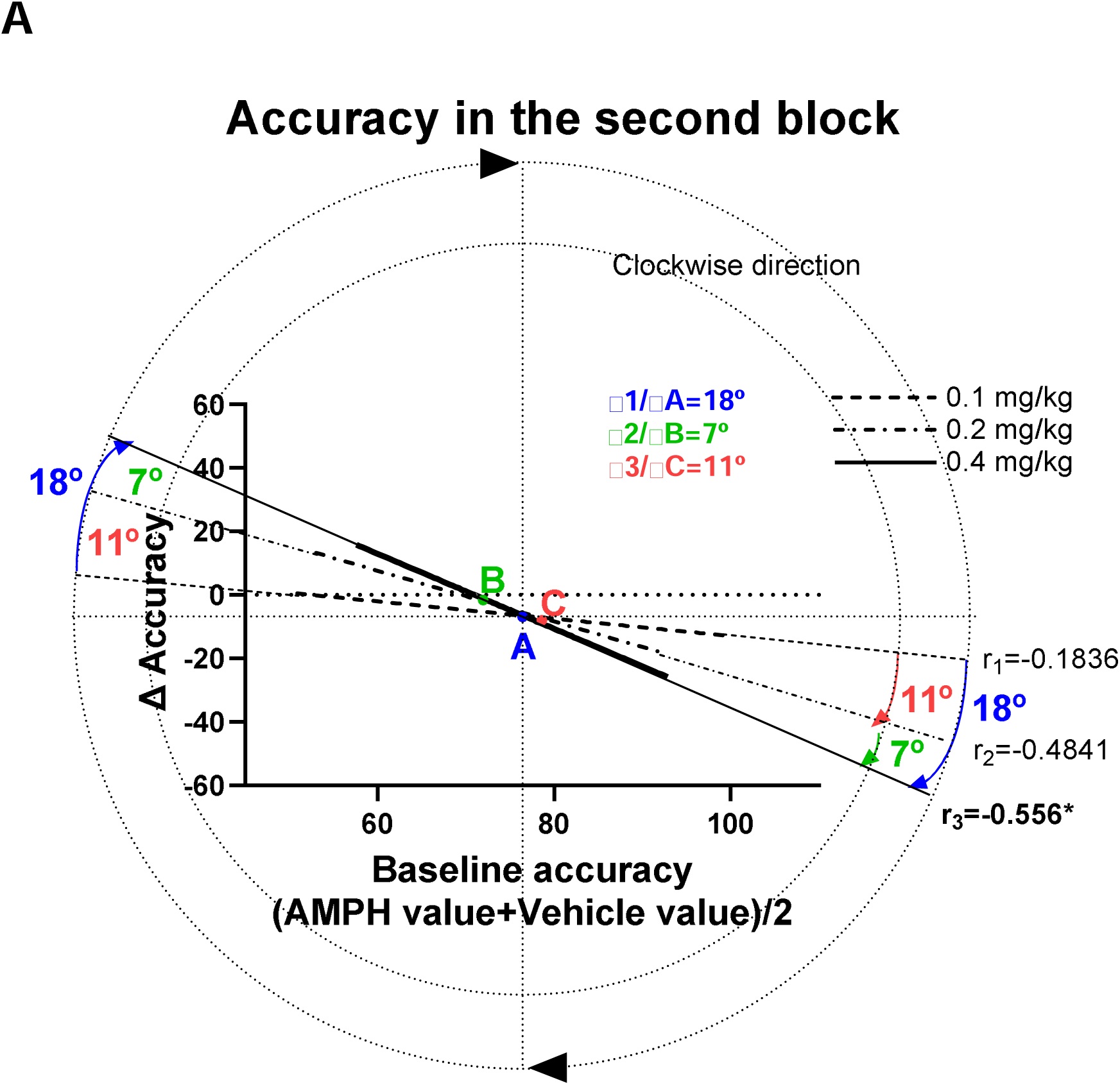
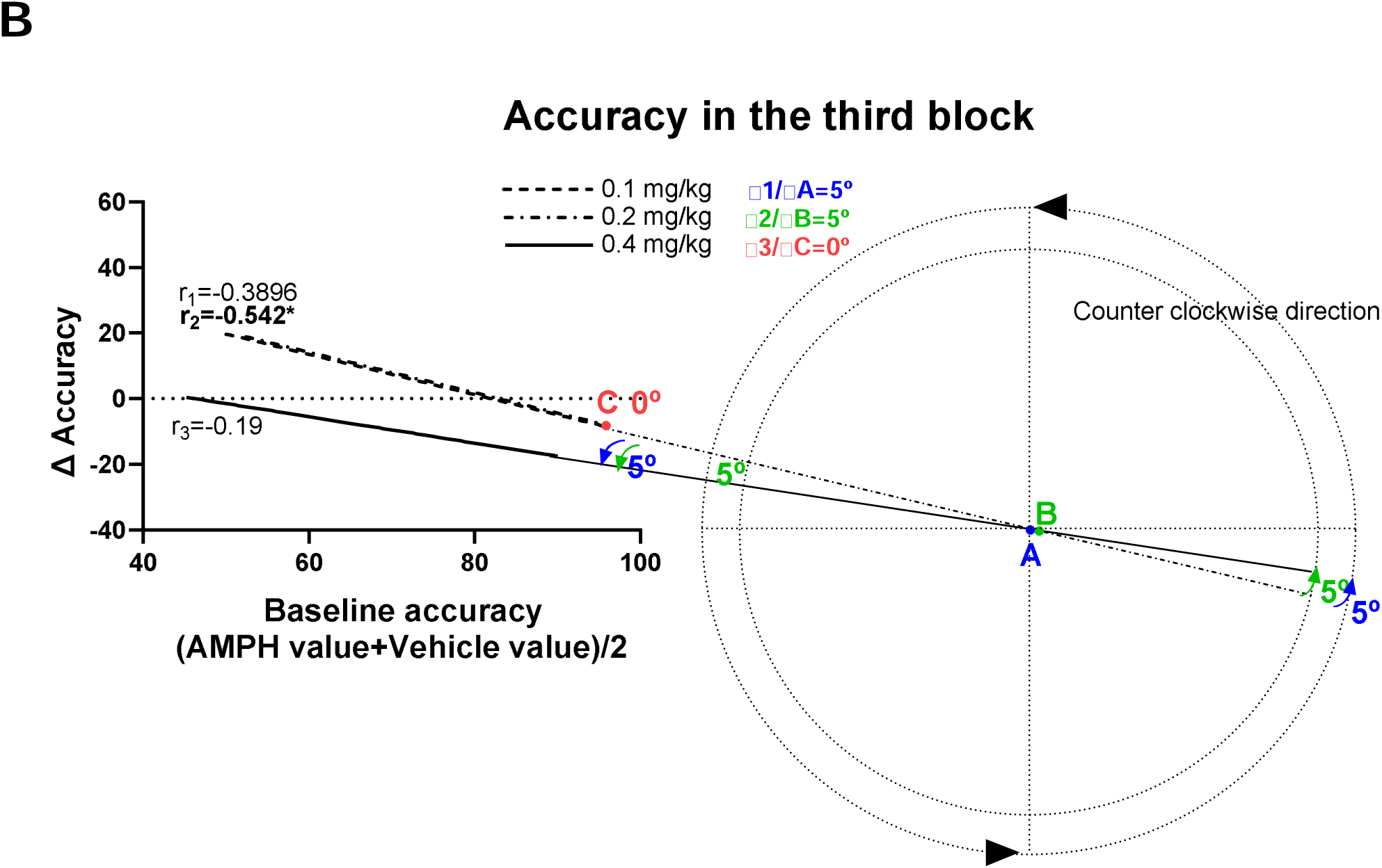
Measurements of Oldham’s angles for accuracy: (A) Accuracy in the second block. (B) Accuracy in the third block.

## Measurements of Oldham’s angles for sensitivity index, hit rate and false alarm rate

Angle measurements for sensitivity index, hit rate and false alarm rate are shown in Fig 25. Amount of space between the regression lines were measured using a protractor. Three angles were measured: ∠1 (blue), ∠2 (green) and ∠3 (red) as shown in Fig 25. Oldham’s angle measurements for the relationship between baseline sensitivity index and AMPH induced change are shown in Fig 25A. ∠1 is the space between the highest and the lowest dose (0.4 mg/kg and 0.1 mg/kg regression lines). ∠2 is the space between the highest and the intermediate dose (0.4 mg/kg and 0.2 mg/kg regression lines). ∠3 is the space between 0.2 mg/kg and 0.1 mg/kg lines (Fig 25). The angle measurement and the direction of movement (clockwise or counterclockwise) shows how effective is the highest dose at increasing sensitivity index and reducing response inhibition of rats compared with the lowest or the intermediate dose. The Exterior Angle theorem states that the exterior angle of a triangle is equal to the sum of the two opposite interior angles. If two angles are known, then the exterior angle theorem can be used to calculate the third angle (Fig 25 B and C).

**Figure 25:**
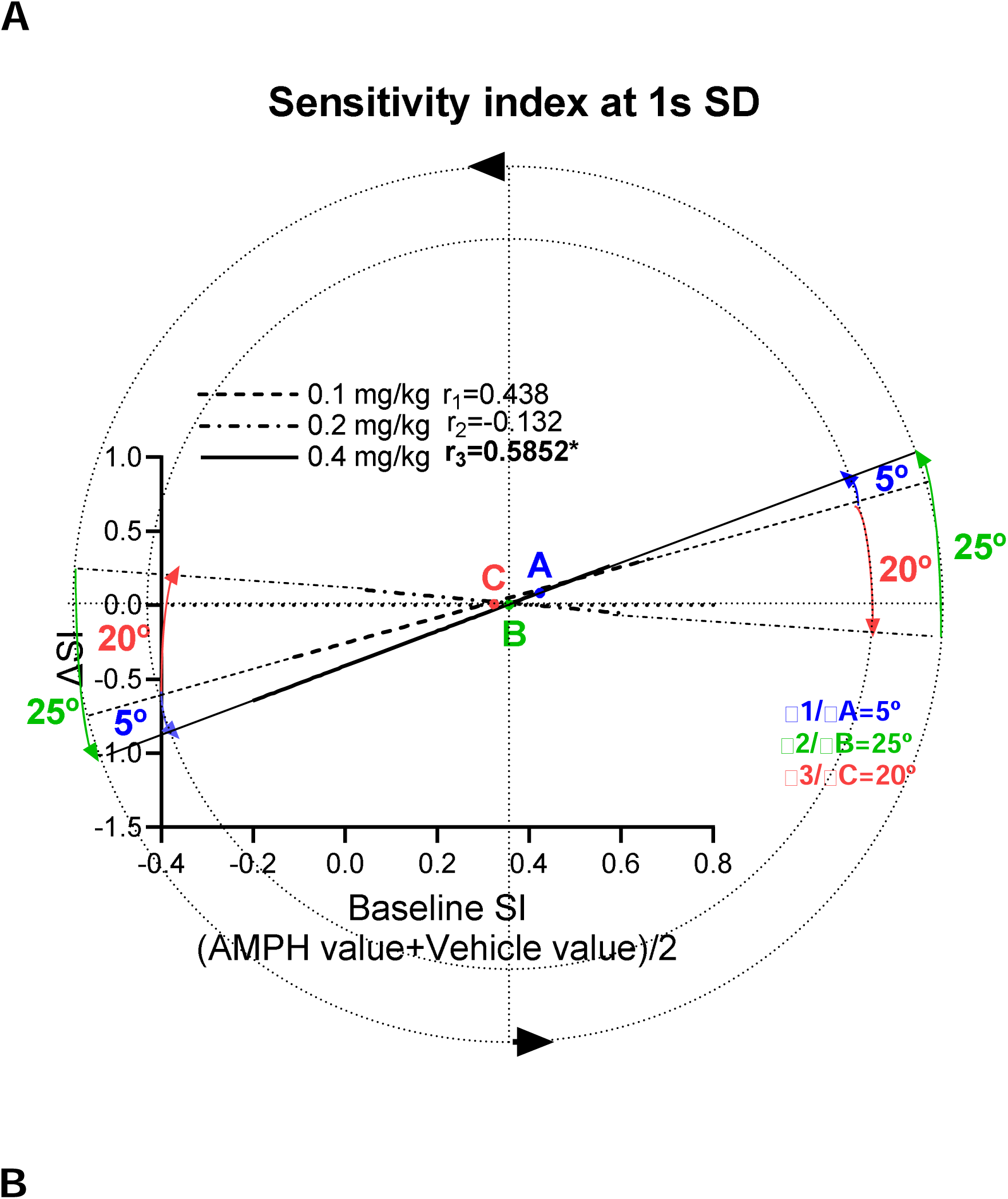
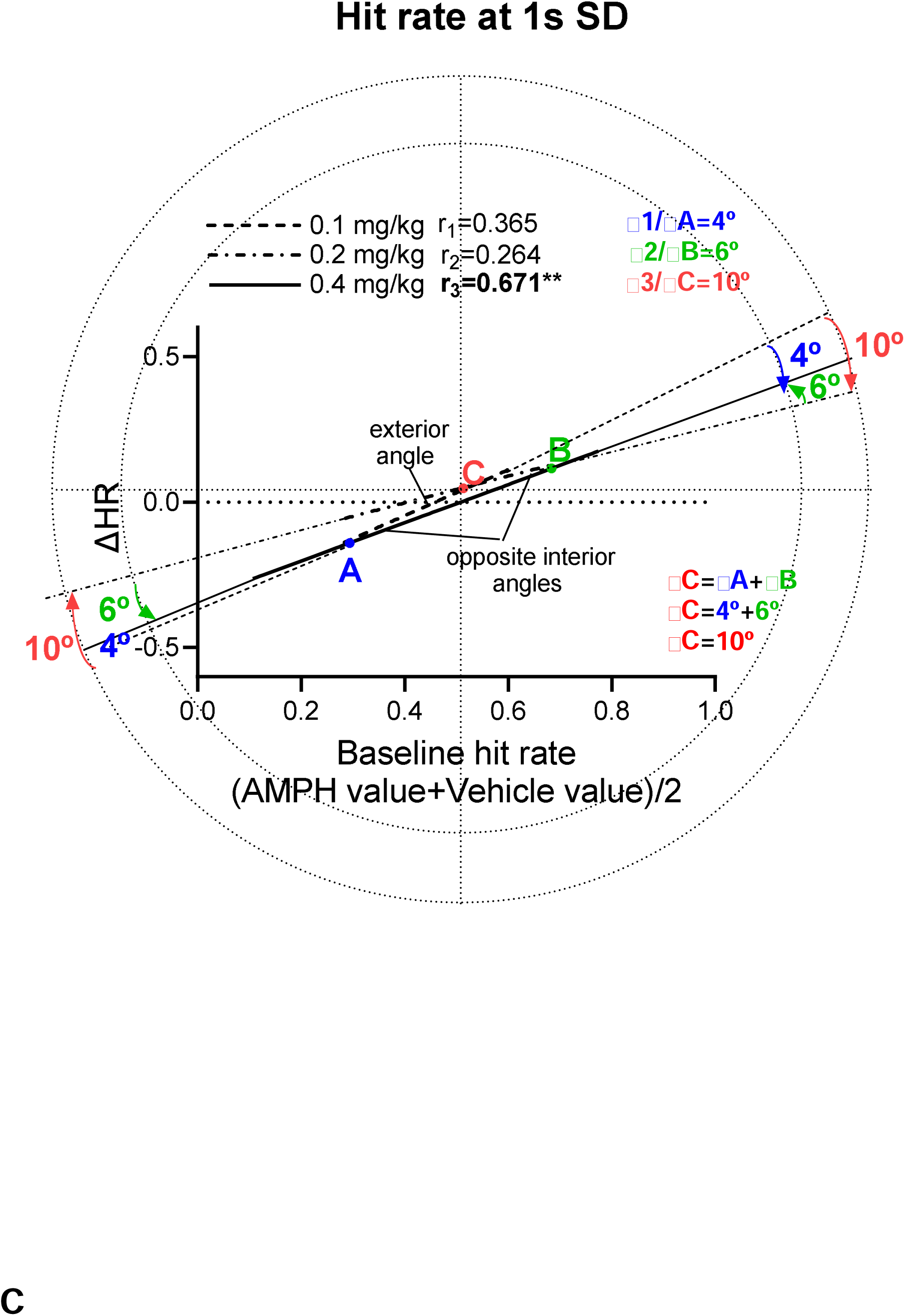
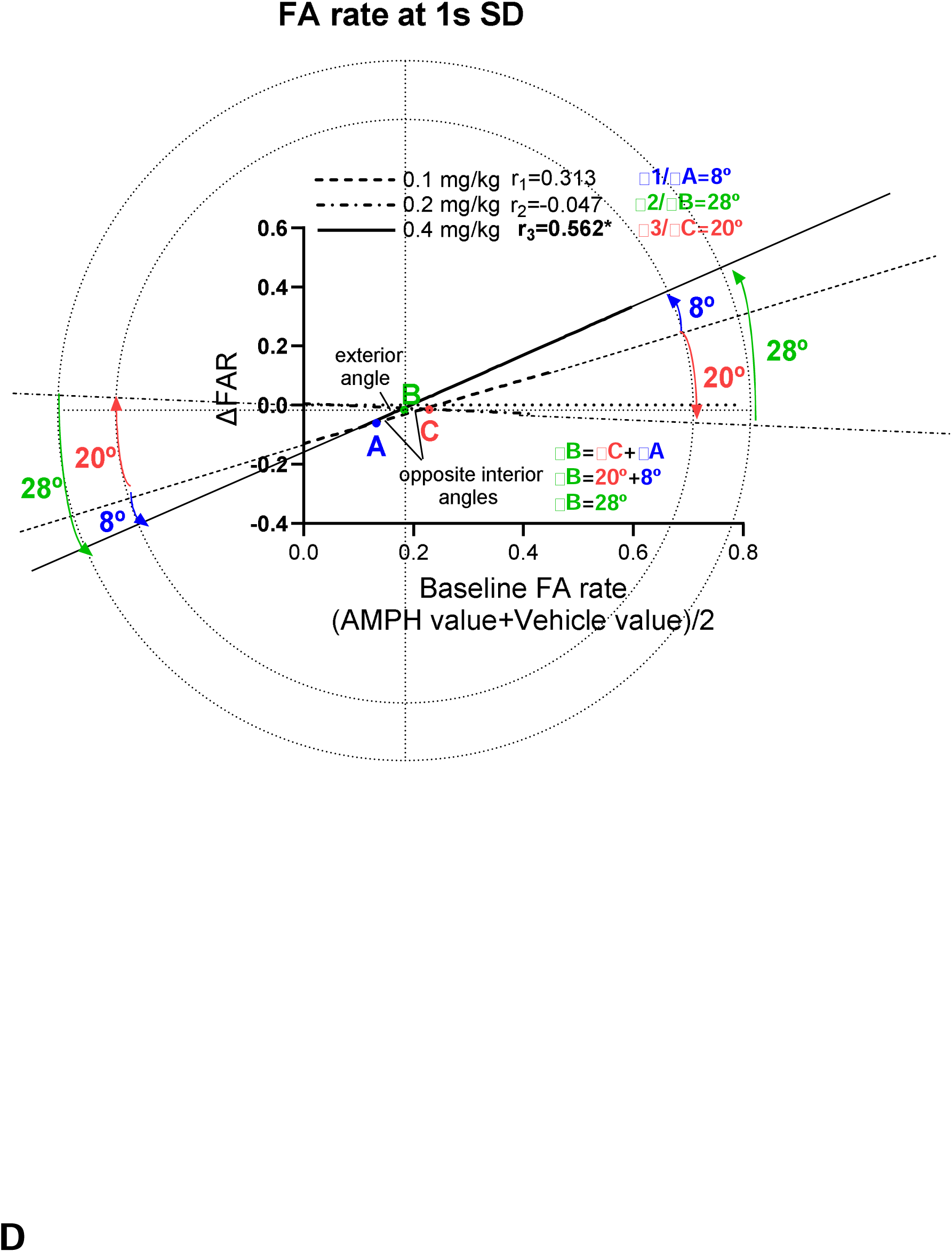
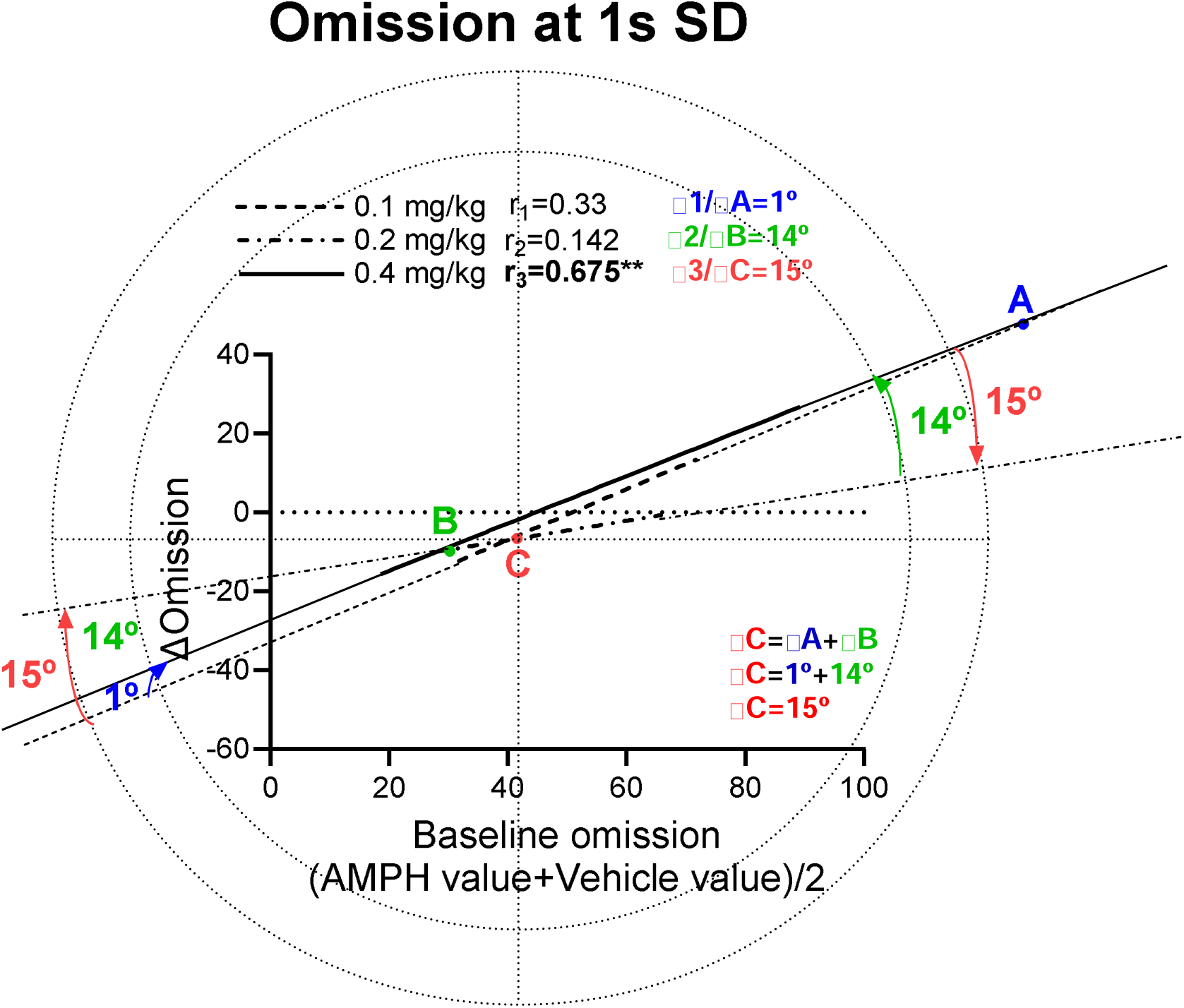
Measurements of Oldham’s angles for: (A) Sensitivity index, (B) Hit rate, (C) false alarm rate and (D) Omission.

Measurements of Oldham’s angles for accuracy at 1.5 second stimulus duration is shown in Fig 26. Amount of space between the regression lines were measured using a protractor. Three angles were measured: ∠1 (blue), ∠2 (green) and ∠3 (red) as shown in Fig 26. ∠1 is the space between the highest and the lowest dose (0.4 mg/kg and 0.1 mg/kg regression lines). ∠2 is the space between the highest and the intermediate dose (0.4 mg/kg and 0.2 mg/kg regression lines). ∠3 is the space between 0.2 mg/kg and 0.1 mg/kg lines (Fig 26). The angle measurement and the direction of movement (clockwise or counterclockwise) shows how effective is the highest dose at increasing accuracy compared with the lowest or the intermediate dose.

**Figure 26:**
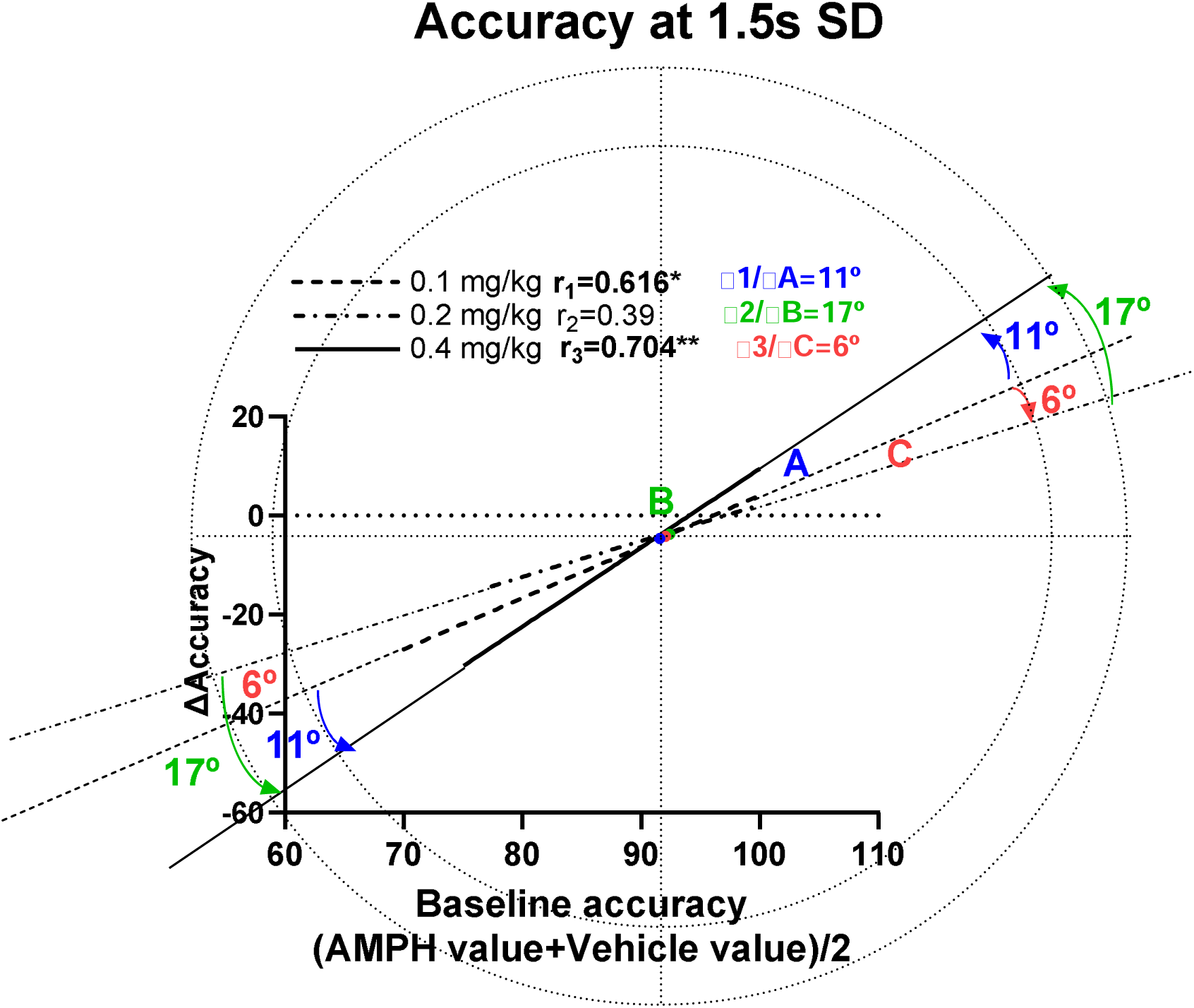
Measurements of Oldham’s angles for accuracy at 1.5 second stimulus duration.

Measurements of Oldham’s angles for sensitivity index are shown in Fig 27. Amount of space between the regression lines were measured using a protractor. Three angles were measured: ∠1 (blue), ∠2 (green) and ∠3 (red) as shown in Fig 27. ∠1 is the space between the highest and the lowest dose (0.4 mg/kg and 0.1 mg/kg regression lines). ∠2 is the space between the highest and the intermediate dose (0.4 mg/kg and 0.2 mg/kg regression lines). ∠3 is the space between 0.2 mg/kg and 0.1 mg/kg lines (Fig 27). The angle measurement and the direction of movement (clockwise or counterclockwise) shows how effective is the highest dose at increasing sensitivity index compared with the lowest or the intermediate dose.

**Figure 27:**
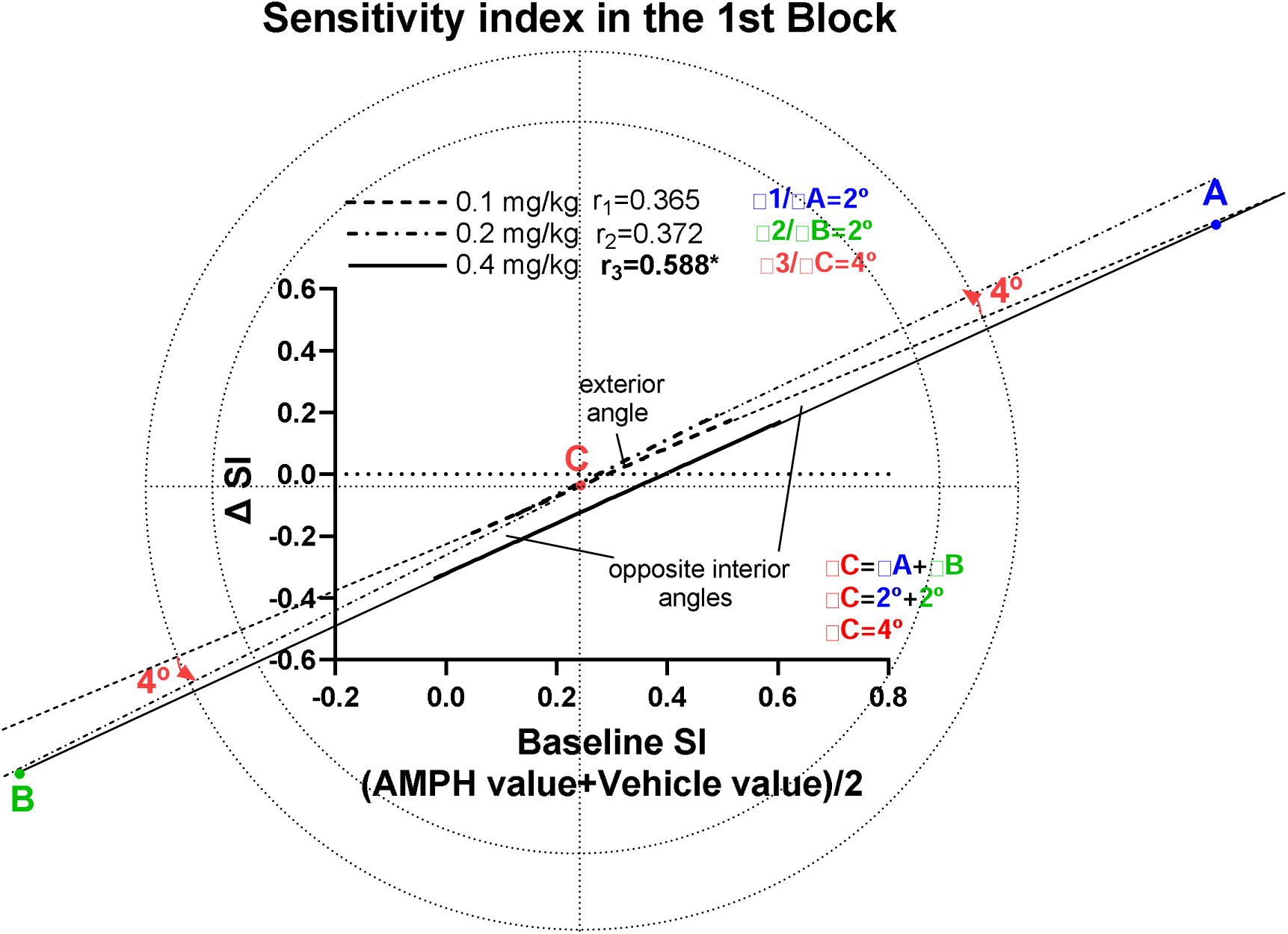
Measurements of Oldham’s angles for sensitivity index.

## Discussion

The present study demonstrates that AMPH impairs attentional processes at the dose of 0.4 mg/kg while lower doses of AMPH show a tendency toward improving attentional performance in a baseline-dependent manner. These results suggest that: (a) the reduced SD challenge changes responsivity index rather than measures of sustained attention (sensitivity index); (b) under this type of challenge AMPH changes attentional performance in a baseline-dependent manner; (c) the type of relationship between baseline performance and the drug-induced change (i.e., inverse or direct relationship) depends on the dose administered; and (d) AMPH changes performance in a dose-dependent manner.

In this study, AMPH disrupted sustained attention in the form of reducing hit rate and increasing false alarm rate under 0.5,1.0,1.5 and 2.0s SDs at the highest dose tested (0.4mg/kg). These effects were caused by a decrease in attentional capacity and not by change in responsivity. Oldham’s method revealed that: (a) the highest dose of AMPH (0.4mg/kg) impaired sustained attention in a baseline dependent manner (direct relationship) at the intermediate SD (1.0s); (b) the highest dose of AMPH impaired accuracy in a baseline dependent manner at 1.5s SD; and (c) the intermediate dose of AMPH (0.2mg/kg) changed accuracy in a rate dependent manner (inverse relationship) at the third trial block. AMPH is known to have beneficial effects on attentional performance in ADHD patients, however, pre-clinical studies to date regarding the effects of AMPH on attentional performance is mixed. Some studies report impairment in attentional accuracy while other studies report improvement in attentional accuracy following administration of AMPH (Turner et al., 2017). This discrepancy could be due to: (1) the task used to measure attention; (2) the challenging condition under which the study was conducted; (3) the species used (rats vs mice); and (4) the doses of the drug administered.

In 5C-CPT studies AMPH has only been tested on mice. For example, in touchscreen 5C-CPT task, 0.3mg/kg dextroamphetamine improved d prime in the form of improving hit rate and accuracy without changing response inhibition. The same dose showed a tendency toward impairing motoric impulsivity (MacQueen et al., 2018). Furthermore, in the mouse rCPT task, amphetamine (0.3, 0.6 and 1mg/kg) improved sustained attention in the form of reducing response inhibition of low-attentive mice without having a significant effect on the high-attentive mice under very short SD (0.2, 0.4, 0.7, 1.1 and 1.8s) (Caballero-Puntiverio et al., 2018). On the other hand, using 5-CSRTT, Fletcher and colleagues (2011) found that their highest dose of AMPH (0.6mg/kg) caused a significant decrease in accuracy in high-attentive rats. This decrease in accuracy was accompanied by a significant increase in waiting impulsivity. In addition, 0.4 and 0.8 mg/kg AMPH significantly decreased accuracy and impaired waiting impulsivity of rats tested under standard task conditions. The work in this study extends previous findings by examining effects of AMPH on attentional performance as measured by the 5C-CPT. The present results, consistent with the impairment in performance reported by previous studies using 5-CSRTT, show that the highest dose of AMPH caused a significant impairment in average SI (decrease in HR+ increase in FAR) at the most challenging SDs (0.5,1 and 1.5s). The disruption in sustained attention following administration of the highest dose of AMPH in this study seems in contrast with efficacy of AMPH in treating ADHD patients. One possible explanation is that the dose used in this study (0.4 mg/kg) was too high. This finding is consistent with the inverted U-shaped relationship between dopaminergic system activation and attentional performance.

Previous findings in 5-CSRTT suggest that reducing SD impairs attentional performance (i.e., accuracy). In this study, reducing SD decreased responsivity to target and non-target trials and increased omission. Interestingly, baseline sensitivity index remained stable for all SDs. This manipulation only affected baseline sensitivity index when the SD was less than 0.5s. However, the mild decrease in baseline sensitivity index at 0.25s SD was due to a significant change in baseline responsivity index. The impairment in attentional processes produced by using a reduced SD challenge does not mimic that of dopamine-compromised rats, as this type of challenge mainly affects responsivity index. The highest dose of AMPH showed a tendency towards reducing attentional measures (accuracy, hit rate and SI) during the intermediate SDs (0.5,1 and 1.5s). The highest dose of AMPH showed a tendency towards increasing FAR at 1.0 and 1.5s SD. The change in sustained attention (SI) at 1.0s SD was caused by change in attentional capacity (decrease in HR and increase in FAR). However, the change in sustained attention at the longer SDs (1.5s or 2s SDs) was caused by change in responsivity.

The standard session length for 5C-CPT is 120 trials while the standard session length for 5-CSRTT is 100 trials. Previous studies report that AMPH does not change accuracy during the standard session length in 5-CSRTT. However, AMPH improved rat 5-CSRTT accuracy when the session length was increased to 250 trials. This improvement in accuracy was observed later in the session, when the control performance declined (Grottick and Higgins, 2002). These findings suggest that extending the session to 250 trials helped in revealing pro-attentional effects of AMPH. It should be taken into consideration that the improvement in attentional accuracy reported in previous studies was observed following administration of low doses of AMPH (0.05 and 0.1 mg/kg) while the highest dose of AMPH (0.4 mg/kg) did not affect accuracy (Grottick and Higgins, 2002). Patients with ADHD cannot maintain their vigilance over time. The present study did not demonstrate vigilance decrement which is decline in vigilance (SI) over time. There was a mild decrease in baseline SI (due to decrease in HR) in the third trial block compared with the first trial block, however, it did not reach a significant level, suggesting an extended session length is needed to demonstrate vigilance decrement. Impairment in sustained attention following administration of 0.4 mg/kg AMPH was observed during the second trial block, mainly due to increase in FAR and, to a lesser extent, decrease in HR. On the other hand, the two lowest doses of AMPH showed a tendency toward increasing SI in the fourth trial block, mainly due to decrease in FAR. Future studies should test lower doses of AMPH (0.05 and 0.1 mg/kg) over extended session length to investigate the pro-attentional effects of this drug on sustained attention.

Correlational analysis was used to account for natural variation in baseline data when evaluating the effectiveness of AMPH on 5C-CPT performance measures. Variation in baseline attentional measures (accuracy, hit rate and SI) was produced by using reduced SDs (0.25, 0.5,1.0, 1.5 and 2s). Oldham’s method is the most appropriate method for determining the presence of a rate dependent relationship (Snider et al., 2016), which would help to identify individuals that would benefit the most from the intervention. Oldham’s method revealed a direct relationship between baseline accuracy and AMPH-induced change (i.e., greater decrease in accuracy in rats with lower baseline accuracy) at the dose of 0.4mg/kg. This type of relationship is less common than the inverse relationship between baseline performance and the drug induced change, suggesting that baseline levels of accuracy can predict magnitude of change in accuracy following administration of AMPH. AMPH did not change other performance measures in a baseline-dependent manner. Although correlational analysis under all SDs revealed no rate-dependent relationship between baseline sustained attention and AMPH induced change, correlational analysis under each SD helped in revealing rate-dependent effects of AMPH on sustained attention. Oldham’s method revealed a direct relationship between measures of attentional performance (SI, HR, FAR, omission) and the change following administration of the highest dose of AMPH during the intermediate SD (1s). This method also revealed the same type of relationship between accuracy and the change following 0.4mg/kg AMPH at longer SD (1.5s). This finding is consistent with previous studies which report the presence of a direct relationship between baseline response rate and change of punished responding for food in pigeon following administration of AMPH (0.1-1.0 mg/kg) (Evenden, 1991) Our data represent the first study to report the rate-dependent effects of AMPH on attentional performance as measured by the 5C-CPT. These findings contribute to the validation of Oldham method in revealing baseline dependent effects of AMPH on 5C-CPT performance measures.

Within-session performance revealed that the highest dose of AMPH impaired SI in a baseline-dependent manner at the beginning of the session. Correlational analysis between Δ performance measures revealed that the Δ SI following administration of 0.4 mg/kg AMPH was due to Δ FAR. However, the baseline-dependent effects of AMPH on SI waned as the session progressed. To our knowledge this is the first study to report baseline-dependent effects of AMPH on measures of sustained attention. As the session progressed, an inverse relationship started to appear between baseline accuracy and the change following administration of 0.2 mg/kg AMPH at the second trial block and it reached a significant level at the 3^rd^ trial block, indicative of a rate-dependent effect. This finding is consistent with previous studies suggesting that selective attention and sustained attention are two different forms of attention and recruit different brain networks. To our knowledge this is the first study to report the presence of an inverse relationship between accuracy as measured by 5C-CPT task and the change following administration of low doses of AMPH, further supporting the inverted U-shaped relationship between attentional accuracy and dopaminergic system activation in 5C-CPT task (Arnsten, 1998; Granon et al., 2000). Furthermore, the present study is consistent with pro-attentional effects of AMPH on selective attention in the signal detection task. In the latter, low doses of AMPH (0.1mg/kg) improved accuracy of low-attentive rats in a rate-dependent manner when they were challenged under reduced stimulus durations (0.06, 0.12, 0.25, 0.5 and1s) while higher doses of AMPH (0.25 mg/kg) did not improve accuracy in a rate-dependent manner (Turner et al., 2017), suggesting that improvement in attentional accuracy depends on the dose. Together, these findings provide preclinical support for the rate-dependent effects of AMPH on 5C-CPT task, previously reported in clinical studies.

In conclusion, the present experiment demonstrated the rate-dependent effects of AMPH on 5C-CPT performance measures under reduced SD challenge. To our knowledge, this is the first successful demonstration of rate-dependent effects of AMPH in the 5C-CPT. The findings of this study further support the potential value of Oldham’s method in revealing baseline-dependent effects of drugs on 5C-CPT performance measures. The present experiment also demonstrated effects of reduced SD challenge on 5C-CPT performance measures. This type of challenge affects responsivity index. Therefore, reduced SD challenge is not the appropriate method for challenging sustained attention.

